# The IFN I response in tumor cells is shaped by PARP7–p300/CBP interactions through distinct loss- and gain-of-function mechanisms

**DOI:** 10.1101/2025.06.30.662483

**Authors:** Ivan Rodriguez Siordia, Sonja Rieth, Rory Morgan, Devon Jeltema, Jonathan Tullis, Jan Orth, Nan Yan, Andreas Marx, Michael Cohen

## Abstract

PARP7, a mono-ADP-ribosyl (MAR) transferase, is a key suppressor of the type I interferon (IFN-I) IFNβ in various tumor cells and a validated drug target. This negative regulation is reversed by small-molecule inhibitors of PARP7 catalytic activity, resulting in increased IFN-β expression. Yet, the mechanism of action of PARP7 inhibitors remains unclear because the relevant substrates of PARP7-mediated MARylation are unknown. Using an optimized analog-sensitive chemical genetic (ASCG) approach, we identified the co-activators, p300 and CBP, as nuclear PARP7 substrates. We identified an α-helical domain in PARP7 essential for p300/CBP interaction, MARylation, and proteasome degradation. Disrupting PARP7–p300/CBP interaction prevents PARP7’s suppression of IFNβ in colorectal cancer cells. p300/CBP reciprocally regulate PARP7’s activity and nuclear localization. Intriguingly, treatment with PARP7 inhibitors increased IFNβ expression more than PARP7 knockout in a p300/CBP-dependent manner. Our findings suggest that in some contexts, IFNβ induction by PARP7 inhibitors occurs via two mechanisms: inhibiting MARylation of p300/CBP (loss-of-function) and stabilizing the PARP7– p300/CBP complex (gain-of-function).

**Teaser:** Chemical genetics discovery of p300 and CBP as substrates of PARP7 that are essential for PARP7-mediated regulation of IFNβ via a dual mechanism.

## Introduction

The cGAS-STING (cyclic GMP-AMP Synthase-STimulator of Interferon Genes) pathway is a key nucleic acid sensor that detects cytosolic viral double-stranded DNA (dsDNA) and activates an innate immune response to defend against infection (*1*). This innate immune response is driven by the induction of type I interferons (IFN-Is), particularly IFNβ. IFNβ is also critical for the induction of adaptive immunity, notably the activation of cytotoxic T cells (CTLs) (*2*). In tumor cells, the cGAS-STING pathway has emerged as a critical mediator between genome instability and innate immune activation (*3*). Disruptions in chromosome segregation during mitosis gives rise to cytosolic dsDNA accumulation which−similar to viral dsDNA− activates the cGAS-STING pathway (*4*). Unsurprisingly, tumor cells use multifarious strategies to suppress IFN-I signaling, thereby evading both intrinsic and cytotoxic T cell–mediated killing. (*5*). Countering these immune evasion tactics is a promising therapeutic strategy in cancer.

PARP7 is an enzyme that belongs to a family of ADP-ribosyltransferases (PARP1–16 in humans), and has emerged as a key suppressor of IFN-I signaling in tumor cells (*6–8*). PARPs transfer ADP-ribose (ADPr) from nicotinamide adenine dinucleotide (NAD^+^) to amino acid residues on protein substrates, a post-translational modification known as ADP-ribosylation (*9*). ADP-ribosylation, similar to ubiquitylation, exists in two forms: mono-ADP-ribosylation or MARylation, which involves the transfer of a single ADPr, and poly-ADP-ribosylation (PARylation), where chains of ADP-ribose are attached to a substrate protein (*10*). Like most PARP family members, PARP7 is thought to catalyze MARylation exclusively (*11*).

Although studies using NAD^+^-competitive inhibitors of PARP7 show that PARP7’s catalytic activity is crucial for suppressing IFN-I expression (*7, 12*), the exact mechanism by which PARP7-mediated MARylation exerts this effect remains unclear. One study proposed that PARP7 MARylates TBK1 (TANK-binding kinase 1), inhibiting its kinase activity (*6*). TBK1 functions downstream of cGAS-STING signaling and is essential for activating the transcription factor IRF3 (Interferon Regulatory Factor 3) through phosphorylation-induced dimerization, which leads to its nuclear translocation and the induction of IFNβ expression (*13*). Accordingly, MARylation of TBK1 by PARP7 is expected to obstruct the phosphorylation and activation of IRF3. Yet, subsequent independent studies indicate that PARP7 inhibitors do not have a discernible effect on TBK1 activation (*12*), and recent findings demonstrate that PARP7 acts downstream of and independently from TBK1 activation to inhibit IRF3-driven transcription (*8*). This finding aligns with several studies showing that PARP7 regulates the function of transcription factors (*11, 14–18*). In some cases, PARP7 represses the functions of transcription factors such as AHR (Aryl Hydrocarbon Receptor), ERα (Estrogen Receptor alpha), and HIF-1α (Hypoxia-Inducible Factor 1-alpha) (*11, 14, 15*). In other instances, PARP7 enhances the functions of transcription factors like LXR (Liver X Receptor), AR (Androgen Receptor), and FRA1 (Fos-Related Antigen-1) (*16–18*). These findings reinforce the emerging view that PARP7’s catalytic activity is critical in transcriptional regulation. However, the mechanism by which PARP7-mediated MARylation regulates IRF3-mediated transcription remains unclear.

Establishing a functional role for PARP7-mediated MARylation—or that of any PARP family member—requires identifying their specific substrates, a task complicated by their shared dependence on NAD⁺ as a substrate. To overcome this, we previously developed an analog-sensitive chemical genetic (ASCG) strategy tailored for PARP7. This involved engineering an analog-sensitive (as)-PARP7 variant capable of utilizing a modified NAD⁺ analog bearing an alkyne handle (*19*). The alkyne can be coupled to biotin using copper-catalyzed azide-alkyne cycloaddition (CuAAC) (“click” chemistry) (*20*). Allowing for the detection and enrichment of MARylated substrates, as demonstrated in similar strategies for other PARPs (*19, 21*). Our ASCG approach for PARP7 analogs successfully identified a few substrates aligned with the known biology of PARP7, as supported by knockout and inhibitor studies (*19*). However, the most significantly enriched substrates we identified were abundantly expressed in HEK 293T cells, raising concerns that our strategy might have overlooked lower-abundance substrates with critical functional roles in regulating IFN-I expression. Based on our unpublished studies, we concluded that the underlying issue stemmed from the CuAAC step, resulting in inconsistent and low stoichiometric labeling. Addressing this obstacle is vital for understanding how PARP7-mediated MARylation regulates IRF3-mediated transcription.

Herein, we describe a third-generation ASCG approach using an NAD^+^ analog directly conjugated to desthiobiotin (DTB), thereby circumventing the need for CuAAC. With this optimized strategy, we identified hundreds of low-abundance nuclear proteins. Intriguingly, key transcriptional regulators, including co-activators and co-repressors, emerge as the most significantly enriched substrates. Notably, for the first time, we demonstrate that p300 (E1A- binding protein p300) and CBP (CREB-binding protein), critical co-activators of IRF3-mediated transcription, are primary PARP7 substrates.

We identified a previously uncharacterized region of PARP7 that is essential for its interaction with and MARylation of p300/CBP. Our data demonstrate that PARP7 MARylation of p300 regulates its stability through a proteasome-dependent mechanism. Intriguingly, we discovered that PROTAC-mediated degradation of p300/CBP reduces PARP7 auto-MARylation, revealing a bidirectional regulatory relationship between PARP7 and p300/CBP. Functional studies show that the interaction between PARP7 and p300/CBP is required for PARP7- dependent suppression of IRF3-driven transcription of IFNβ in the mouse colon carcinoma cell line, CT26.

Unexpectedly, pharmacologic inhibition of PARP7 using the small-molecule PARP7 inhibitor RBN2397 increased IFNβ levels beyond those observed in PARP7 knockout CT26 cells—an effect dependent on the PARP7–p300/CBP interaction. Furthermore, we found that RBN2397 and the structurally distinct PARP7 inhibitor KMR206 induce hyperphosphorylation of p300/CBP in the context of cGAS/STING activation.

Collectively, our findings reveal a complex regulatory network between PARP7 and p300/CBP that is critical for suppressing IRF3-driven transcription. Our data also uncover a previously unappreciated “dominant positive” effect of PARP7 inhibitors, which has important implications for the therapeutic development of PARP7 inhibitors.

## Results

### Third-generation ASCG strategy reveals transcriptional co-activators p300 and CBP as primary substrates of PARP7

In the cGAS-STING pathway, TBK1 is the key kinase that drives IRF3 activation through phosphorylation-induced dimerization and nuclear translocation. While TBK1 was initially proposed as the direct substrate of PARP7 MARylation, leading to the regulation of IRF3- mediated transcription and downstream IFNβ signaling (*6*), recent evidence indicates that PARP7 instead acts downstream of IRF3 activation (*8*). Moreover, in many cell types, PARP7 is predominantly localized in the nucleus (*12, 18*), raising questions about how it could MARylate a cytoplasmic protein like TBK1.

In earlier studies, we established an analog-sensitive chemical genetic (ASCG) strategy for the unbiased identification of direct PARP7 substrates (*19*). In this approach, we engineered an analog-sensitive PARP7 (I631G PARP7, IG-PARP7) to uniquely utilize an NAD^+^ analog modified at the C-5 position of the nicotinamide ring with a benzyl (Bn) group that prevents binding to WT-PARPs. This 5-Bn-NAD^+^ analog also features an alkyne group, which serves as a latent handle for coupling to biotin via “click” chemistry. Biotin conjugation enables identification of direct PARP7 substrates using avidin-based enrichment followed by tandem mass spectrometry. Our first-generation ASCG approach featured a 5-Bn-NAD^+^ analog with an alkyne (propargyl) at the *N-6* position (Fig. 1A, Top), but this analog was an inefficient substrate of IG-PARP7 (*19*). Using structure-guided design, we generated a 5-Bn-NAD^+^ analog containing an alkyne (ethynyl) at the C-2 position of the adenine ring (Fig. 1A, Middle). Our second-generation ASCG approach using the C-2 ethynyl 5-Bn-NAD^+^ successfully identified direct substrates of PARP7, several of which aligned with the known biology of PARP7. However, the most significantly enriched substrates we identified were abundantly expressed in HEK 293T cells, raising concerns that our strategy might have overlooked lower-abundance substrates with critical functional roles. We hypothesized that the underlying issue stemmed from the CuAAC step. For instance, the efficacy of CuAAC-mediated labeling of as-PARP substrates proved highly sensitive to protein concentration in the lysates. Moreover, due to the low stoichiometry of analog-sensitive PARP-mediated MARylation with our clickable 5-Bn-NAD^+^ analogs (true also for native NAD^+^ with WT-PARPs), the excess biotin azide compromised the enrichment of as- PARP substrates. Overcoming the current limitations in our ASCG approach is crucial for uncovering substrates that will provide mechanistic insight into how PARP7-mediated MARylation regulates transcription.

**Fig. 1.**
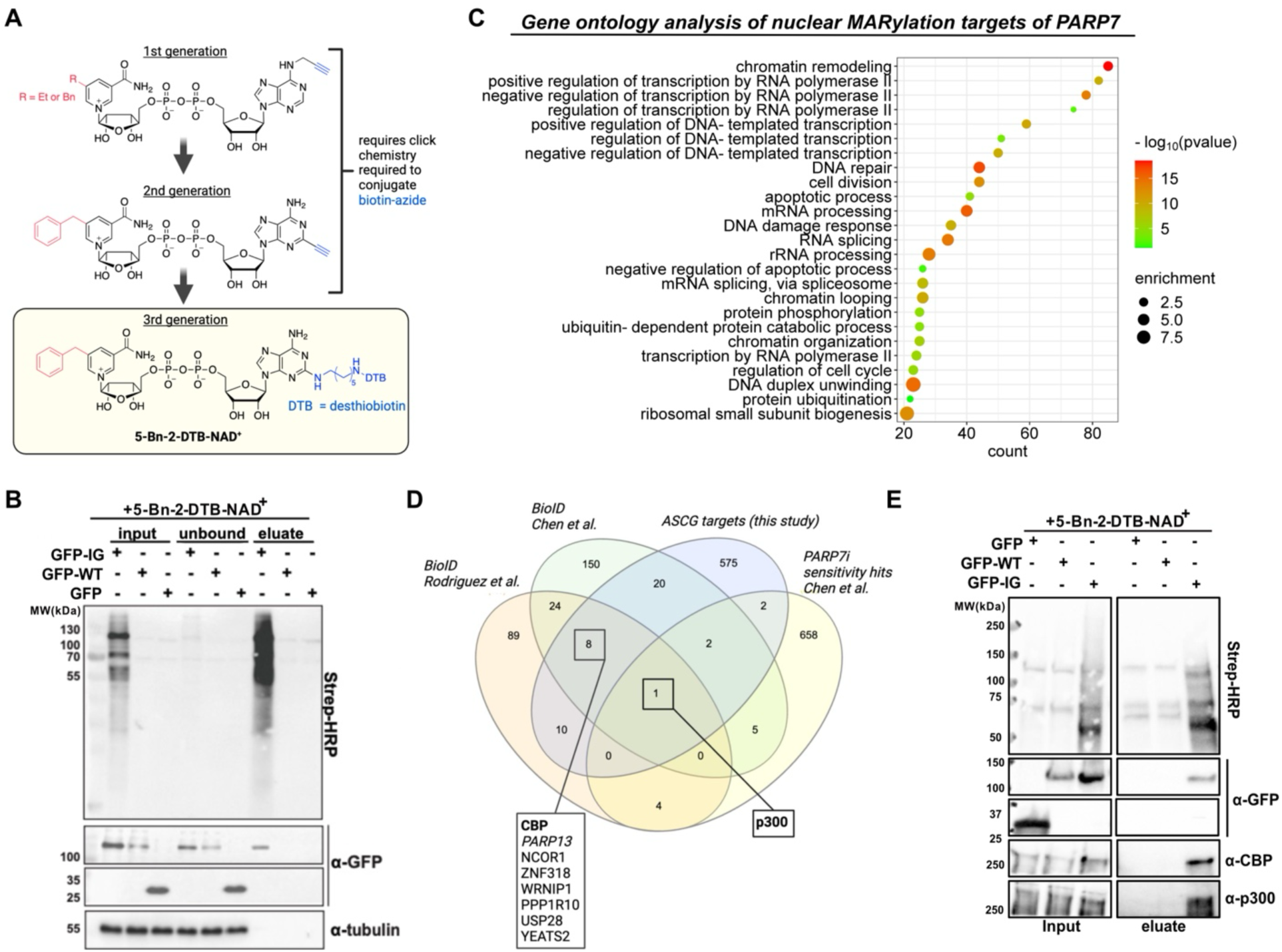
Third-generation ASCG strategy reveals transcriptional co-activators p300 and CBP as primary substrates of PARP7. (**A**) Chemical structure of the NAD^+^ analogs used in the chemical-genetic approach to identify PARP7 targets in HEK 293T cell lysate. Top: 5-Bn-2-e-NAD^+^ in combination with copper(I)- catalyzed azide-alkyne cycloaddition (CuAAC) for functionalization used in Rodriguez et al. Bottom: optimized NAD^+^ analog, 5-Bn-DTB-NAD^+^, containing the alkyne group on C2 of the adenine moiety was replaced by the affinity tag DTB. (**B**) Western blot of HEK 293T cells expressing GFP-PARP7-LG labeled with 100 µM 5-Bn- DTB-NAD⁺ at 30 °C for 2 h. Proteins were precipitated with MeOH/CHCl3 and resuspended in 6 M Urea in 1X PBS with 0.5% SDS. The modified proteins were detected by immunoblotting with ExtrAvidin® peroxidase (anti-Avidin), which binds the DTB tag. Samples were collected after the ADP-ribosylation reaction (input), after incubation with High-Capacity NeutrAvidin™ Agarose beads (unbound), and after the elution of proteins. Controls were performed using cell lysates expressing WT-PARP7 or EV-GFP. (**C**) Gene ontology (GO) analysis of the nuclear substrates of PARP7. Analysis was performed using DAVID. (**D**) Venn diagram showing PARP7 interactors, substrates, and sensitive genes identified in this study, Rodriguez et al., and Chen et al. (**E**) Western blot detecting PARP7-dependent MARylation of p300 and CBP in nuclei isolated from GFP-PARP7-LG–expressing HEK 293T cells. Samples were labeled and processed as in (B).

To address the challenges associated with CuAAC, we established a third-generation ASCG approach, synthesizing a 5-Bn-NAD^+^ analog containing desthiobiotin (DTB) at the C-2 position of adenine (5-Bn-2-DTB-NAD^+^) (Fig. 1A, Bottom, and see Materials and Methods). We chose DTB rather than biotin because it has a lower affinity for streptavidin and related biotin-binding proteins than biotin, allowing for more efficient elution of DTB-conjugated proteins (*22*). DTB conjugates of NAD^+^ and our 5-Et-NAD^+^ analog (our substrate for analog-sensitive PARP1) proved effective for enriching and identifying PARP1 substrates (*23*).

We first determined whether 5-Bn-2-DTB-NAD^+^ is a preferred substrate for IG-PARP7, which can be used to label and efficiently enrich direct substrates of PARP7. We transiently expressed GFP alone, GFP-tagged IG-PARP7, or WT-PARP7 in HEK 293 cells. Cell lysates were prepared and incubated with 5-Bn-2-DTB-NAD^+^ (100 μM). DTB-modified (i.e., MARylated) were enriched using high-capacity NeutrAvidin™ agarose and eluted under denaturing conditions in the presence of free biotin. Consistent with previous studies, IG-PARP7 efficiently used 5-Bn-2-DTB-NAD^+^ for auto- and trans- MARylation as evidenced by DTB labeling observed in western blot analysis (Fig. 1B, lane 1, ‘input’). In contrast, we did not detect DTB labeling in lysates derived from WT-PARP7- or GFP-expressing cells, indicating that 5-Bn- 2-DTB-NAD^+^ cannot be used as a substrate for WT-PARP7 (Fig 1B, lanes 2 and 3). By comparing the ‘input’ (lane 1) with the ‘unbound’ (lane 4) and ‘eluate’ (lane 7) samples from IG- PARP7-expressing lysates, it is evident that DTB-labeled proteins were efficiently enriched and recovered (Fig. 1B).

With 5-Bn-2-DTB-NAD^+^ established as an ideal orthogonal substrate for IG-PARP7, capable of labeling and enriching the direct substrates of PARP7, we next sought to identify these substrates using liquid chromatography-tandem mass spectrometry (LC-MS/MS). We scaled up the DTB labeling experiments described and performed them in triplicate. Data-independent acquisition (DIA) identified thousands of proteins, with 1,883 significantly enriched in IG-PARP7 samples compared to WT-PARP7 and GFP samples (ANOVA, FDR = 0.001)." (Table S1).

Approximately 600 direct PARP7 substrates are annotated as nuclear proteins (Table S1), many of which are low-abundance proteins according to OpenCell (https://opencell.sf.czbiohub.org/). Gene ontology analysis (DAVID) revealed that many of the identified nuclear PARP7 substrates were significantly enriched in pathways related to chromatin remodeling and transcriptional regulation (Fig. 1C and Table S1). While some transcription factors, like AHR (aryl hydrocarbon receptor), a previously described substrate of PARP7 (*11*) were identified as direct PARP7 substrates using our third-generation ASCG approach, many of the most enriched substrates are known transcriptional coregulators, including coactivators and corepressors. Together, our proteomic findings suggest that PARP7 plays a role in regulating transcription through MARylation of coactivators and coregulators, in addition to transcription factors themselves.

Notably, for the first time, we demonstrate that the coregulators p300 (E1A-binding protein p300) and CBP (CREB-binding protein) are highly enriched, direct substrates of PARP7. P300 and CBP contain a histone acetyltransferase (HAT) domain, and their primary role in transcriptional regulation is histone acetylation, which is thought to create an “open” chromatin state, allowing for efficient transcription (*24*). However, p300/CBP also has non-catalytic roles in transcriptional regulation (*25, 26*). While they are often viewed as redundant in many cellular contexts, recent studies indicate that they possess distinct functions in cells (*27, 28*). p300/CBP were particularly intriguing as they are well-established co-regulators of the transcription factor IRF3 (*29, 30*), which are essential for robust transcription of *IFNβ*. CBP was previously identified as a PARP7 interactor in two independent protein-proximity (BioID) labeling mass spectrometry studies (Fig. 1D) (*19*). Additionally, p300 emerged as a significant hit in several genome-wide CRISPR screens to identify genes that regulate resistance to the PARP7 inhibitor RBN2397 (Fig. 1D) (*31, 32*). We did not identify p300/CBP as direct substrates of PARP7 using our second-generation ASCG approach, likely because they are not highly expressed (< 100 nM) in HEK 293T cells (OpenCell). The only nuclear PARP7 substrates identified in both ASCG approaches were PARP13, FAM98A, and RBMX, all of which are highly expressed in HEK 293T cells (> 500 nM), along with WRNIP, which we previously identified as a promiscuous PARP substrate. Take together, our third-generation ASCG approach using 5-Bn-2-DTB-NAD^+^ enabled the identification of lower-abundance nuclear PARP7 substrates, like p300/CBP, many of which have known roles in transcriptional regulation.

To further evaluate p300/CBP as substrates of PARP7 in a more physiological context, we conducted our ASCG approach in intact nuclei, where native transcriptional complexes are better preserved compared to lysate conditions. We reasoned that 5-Bn-2-DTB-NAD^+^ could readily pass through the nuclear pore. Using this strategy, we found that PARP7 MARylates p300/CBP in intact nuclei (Fig. 1E). Finally, we confirmed that p300 is directly MARylated by PARP7 *in vitro* using native NAD^+^ and recombinantly expressed proteins (Fig. S1). PARP13, which we previously identified as a PARP7 substrate using our second-generation ASCG approach, was used as a positive control (Fig. S1).

### Modeling using AlphaFold3 points to a novel interaction between PARP7 and p300/CBP

To gain insight into PARP7–p300/CBP interactions, we modeled PARP7 binding to p300 and CBP using AlphaFold 3 (AF3) (*33*). The AF3 model revealed that the most confidently predicted interaction occurred between the three-helix bundle of the KIX domain of p300/CBP and three uncharacterized α-helices in the N-terminal region of PARP7 (Fig. 2A and Fig. S2). We named these three helices (KIX binding regions 1, 2, and 3 (KBR1, 2, and 3). The three-helix bundle KIX domain of p300/CBP binds to the activation domain of several transcription factors, including CREB, MLL, FOXO3a, c-Myb, and E-protein (*34–38*). This is critical for recruiting p300/CBP to the promoters of these transcription factors to acetylate histones. Interestingly, confidence in the predicted structure of the N-terminal helices of PARP7, in the absence of p300/CBP, is low, and the conformation appears distinct (Fig. S2). This is reminiscent of the regions of certain transcription factors, such as MLL, which are intrinsically disordered but fold into α-helices upon binding to the KIX domain (*39, 40*). Comparing solved NMR structures of the KIX domain bound to various transcription factors to the PARP7–p300 AF3 model showed KBR1-3 of PARP7 engage with the three-helix bundle KIX domain in a manner reminiscent of the KIX-binding α-helices of transcription factors (Fig. 2B). Most transcription factors interact with the KIX domain via a single α-helix, binding either the Myb or MLL binding site (Fig. 2B). In contrast, PARP7 is predicted to interact with both sites via the KBR1 and KBR3 α-helices, resembling the binding mode of FOXO3a (Fig. 2B). All KIX-binding α-helices share a ΦXXΦΦ motif (Φ = hydrophobic amino acid and "X" is any residue) (*38*). Multiple sequence alignment analysis of KIX-binding α-helices with KBR1 and KBR3 revealed that KBR1 and KBR3 contain the conserved ΦXXΦΦ motif (Fig. 2C). Analysis using the AF3 tool from PREDICTOMICS (*38*) to assess confidence in each atom/residue pair revealed that KBR3 displays the lowest inter-residue predicted aligned error (PAE) and the strongest predicted protein–protein interaction interface (Fig. 2D and 2E, Fig. S3B, Table S2). Collectively, these modeling studies support the conclusion that previously uncharacterized N-terminal α-helices, KBR1-3, of PARP7 interact with the three-helix bundle of the KIX domain, analogous to KIX-binding α-helices of transcription factors.

**Fig. 2.**
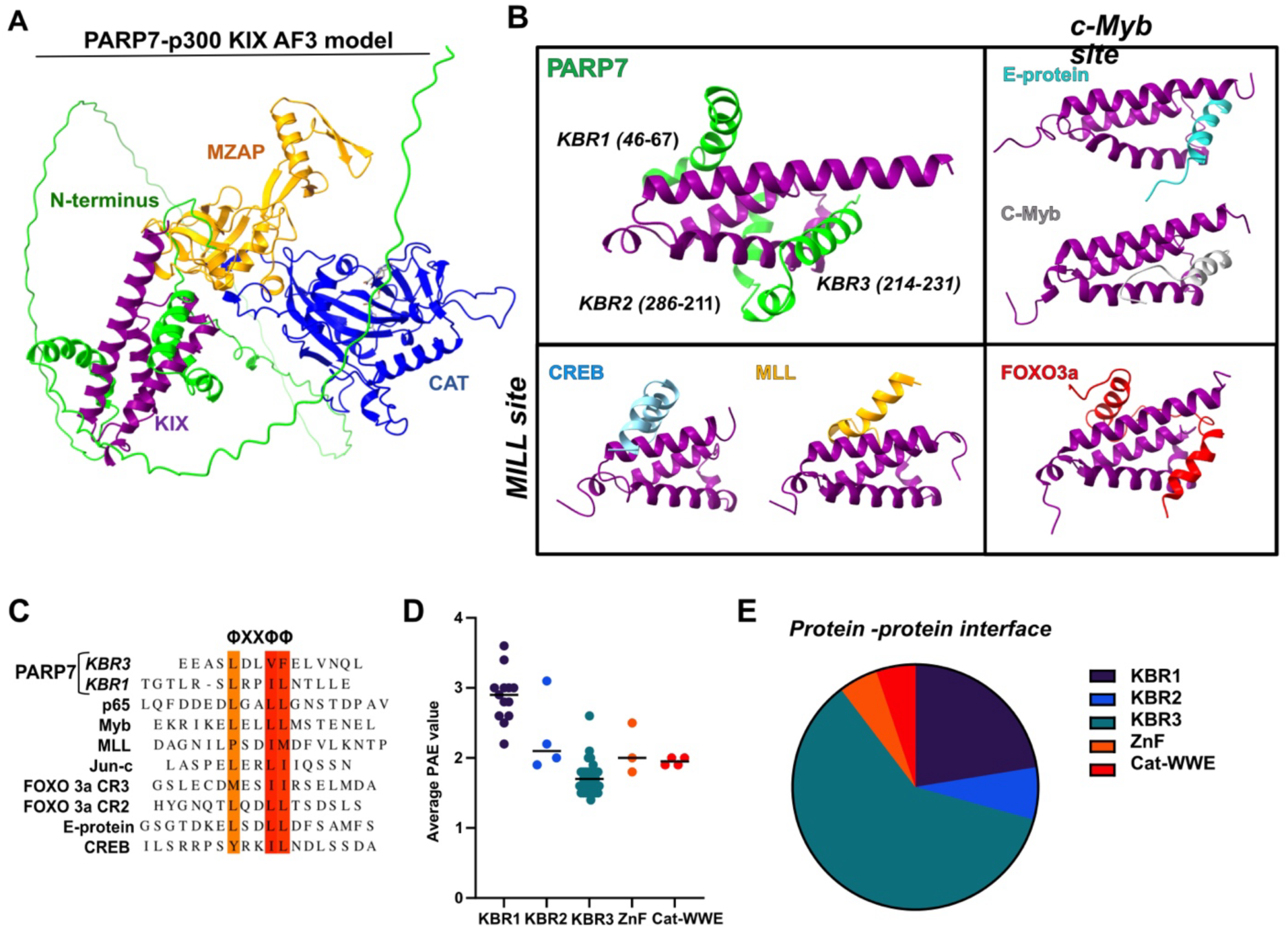
Modeling using AlphaFold3 points to a novel interaction between PARP7 and p300/CBP. (**A**) Simplified AlphaFold 3 prediction of the PARP7-p300 complex. PARP7 N-terminus (green), MZAP (yellow), PARP domain (blue), and p300 KIX domain (purple). The model was created with the full length of p300-PARP7. Non-PARP7 p300 interacting regions were removed for clarity. (**B**) Overlay analysis of the KBR3-KIX complex compared with resolved structures of KIX binding proteins: CREB (blue); MLL (yellow); FOXO3a (red); c-Myb (gray); E-protein (cyan). (**C**) Multiple sequence alignment of KBR1 and KBR3 with other KIX-binding motifs. (**D**) Plot showing predicted aligned error (PAE) assigned in the PARP7-p300 model. (**E**) Pie chart showing PARP7 domains contributing to the p300 interface.

### PARP7 KBR3 is necessary for p300/CBP binding and MARylation

We next sought to evaluate the contribution of KBR3 in the interaction between PARP7 and p300/CBP. We generated a GFP-PARP7 variant in which KBR3 (amino acids 214-231, human numbering) was deleted (ΔKBR3). We assessed p300/CBP binding to PARP7 by co-immunoprecipitation (co-IP) from HEK 293T lysates expressing GFP-tagged WT- or ΔKBR3- PARP7, using GFP-trap beads. Consistent with previous BioID studies, p300/CBP bound to WT- PARP7 (Fig. 3A). Treatment with PARP7 inhibitor RBN2397 (300 nM) (*7*), which increases GFP-WT-PARP7 levels as we and others have observed previously (*18, 19, 41*), did not impact binding of p300/CBP to PARP7 if one considers the increased levels of PARP7 in RBN2397 treated cells (Fig. 3A). Deletion of KBR3 in PARP7 completely abolishes p300/CBP binding to PARP7 (Fig. 3A). Despite comparable increases in the levels of ΔKBR3-PARP7 upon RBN2397 treatment relative to WT-PARP7, we did not observe p300/CBP binding to ΔKBR3-PARP7 (Fig. 3A). These results demonstrate that PARP7 binds to p300/CBP independent of its catalytic activity, and that KBR3 is required for PARP7–p300/CBP interaction.

**Fig. 3.**
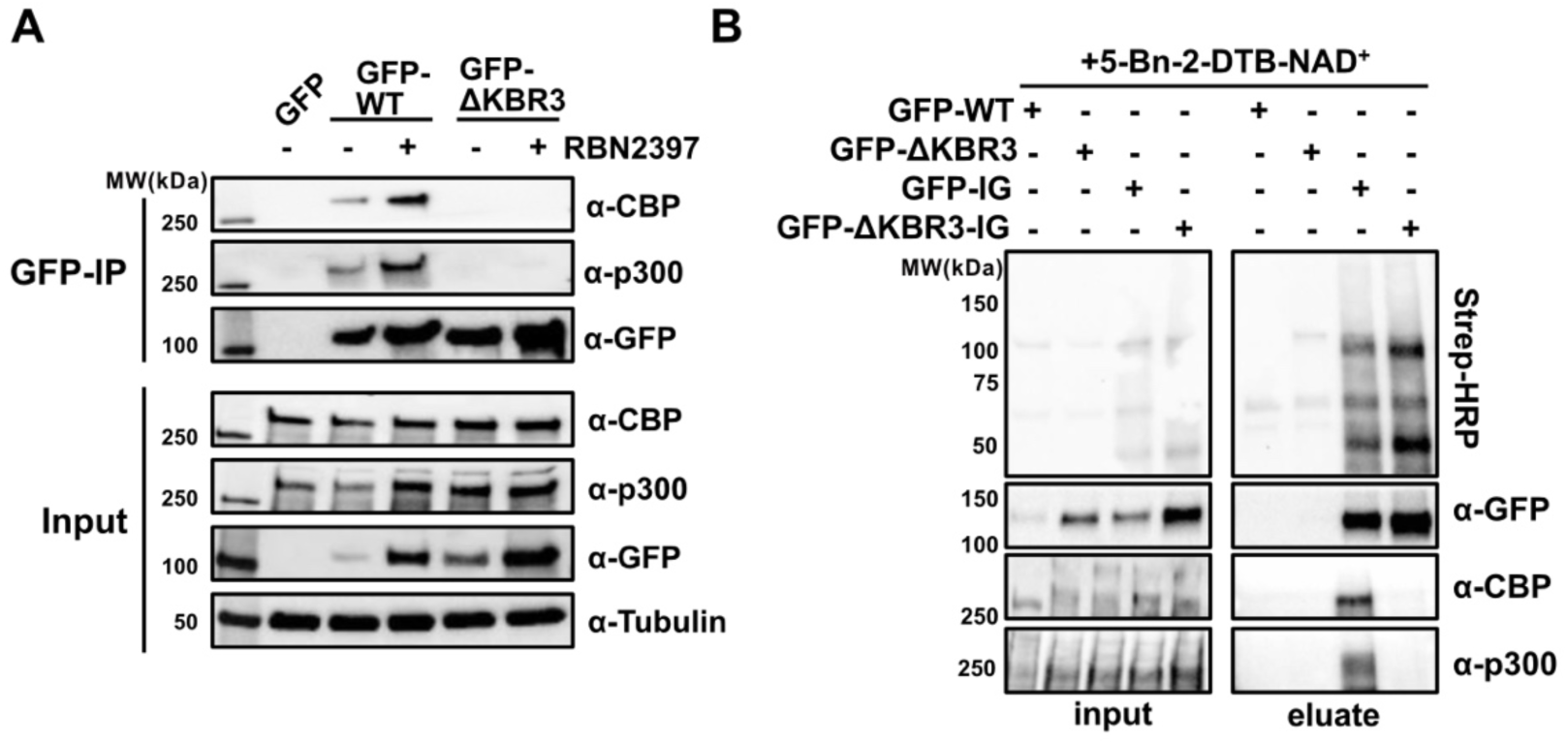
PARP7 KBR3 is necessary for p300/CBP binding and MARylation. (**A**) IP of GFP-PARP7-WT and GFP-PARP7-Δ KBR3 overexpressed in HEK 293T cells +/- 300 nM RBN2397 for 20 h before harvest and IP. (**B**) Western blot showing p300 and CBP MARylation by PARP7-WT versus ΔKBR3. Intact nuclei from HEK 293T cells expressing GFP-PARP7-LG-ΔKBR3 were labeled with 5-Bn-DTB- NAD⁺ and processed as in Fig. 1B. Controls were performed using cell lysates expressing PARP7-WT, PARP7-LG, or PARP7-ΔKBR3.

Given the essential role of KBR3 in mediating the PARP7–p300/CBP interaction, we reasoned that deleting KBR3 would prevent p300/CBP MARylation by PARP7. To test this idea, we generated ΔKBR3 in our analog-sensitive PARP7 variant, IG-PARP7, to directly assess the impact of KBR3 deletion on p300/CBP MARylation in intact nuclei using 5-Bn-2-DTB-NAD^+^. In nuclei derived from cells expressing IG-PARP7, both IG-PARP7 (auto-MARylation) and p300/CBP (trans-MARylation) were present in the NeutrAvidin agarose elutes; in contrast, while ΔKBR3-IG-PARP7 was detected in the elutes—indicating that it is still capable of auto-MARylation—p300/CBP was not (Fig. 3B). These results demonstrate that KBR3 is required for p300/CBP MARylation by PARP7 and further confirm that p300/CBP are bona fide substrates of PARP7.

### PARP7 catalytic activity regulates p300 stability in a proteasome-dependent manner

A closer inspection of the input lanes in the GFP trap co-IP revealed that the expression of WT-PARP7, but not ΔKBR3-PARP7, reduced p300 and, to a lesser extent, CBP protein levels (Fig. 3A). This suggests the intriguing idea that PARP7-mediated MARylation of p300/CBP regulates its stability. The ubiquitin-proteasome system (UPS) is the major regulator of protein stability in cells. To determine if the UPS is involved in PARP7-dependent regulation of p300 levels, we treated HEK 293T cells expressing GFP, GFP-tagged WT PARP7, or ΔKBR3-PARP7 with either bortezomib (100 nM), a potent proteasome inhibitor, or DMSO control. The GFP-WT- PARP7-dependent reduction in p300 levels was prevented by bortezomib (Fig. 4A, B). We observed similar results with RBN2397 (300 nM) treatment, and the combination with bortezomib did not further increase p300 levels (Fig. 4A, B). As anticipated from previous results, ΔKBR3-PARP7 did not influence p300 levels (Fig. 4A, B).

**Fig. 4.**
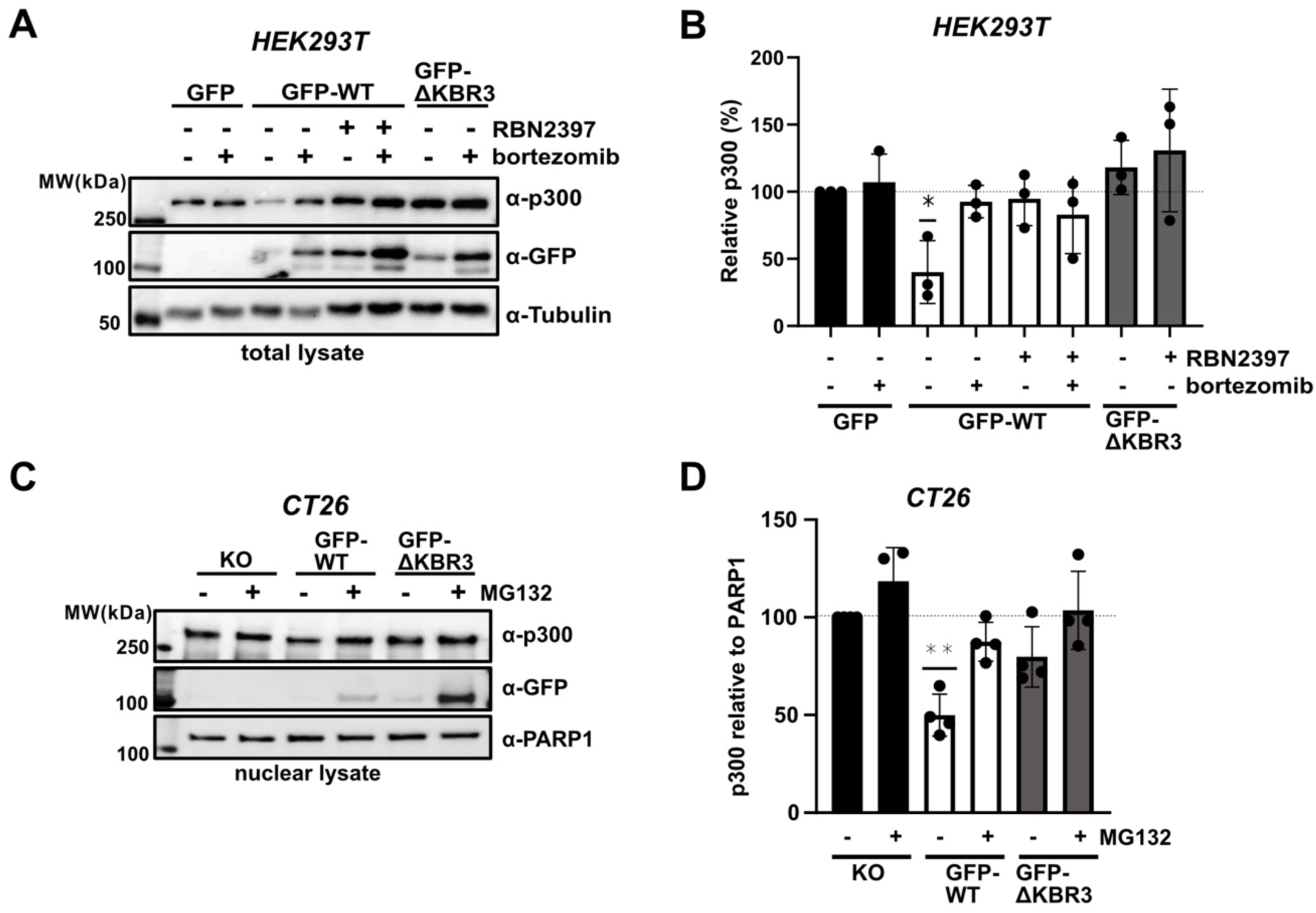
PARP7 catalytic activity regulates p300 stability in a proteasome-dependent manner. (**A**) Western blot of HEK 293T cells expressing GFP-EV, GFP-PARP7-ΔKBR3, and GFP- PARP7-WT ±300 nM RBN2397 for 20 h before treatment with 10 µg/mL DMXAA and 100 nM Bortezomib for 6 h. (**B**) Quantification of three biological replicates from (A). (**C**) Western blot of doxycycline-inducible CT26 cells expressing PARP7-WT or ΔKBR3, treated with 5 µg/mL Dox for 24 h, followed by 5 µM MG132 and 10 µg/mL DMXAA for 6 h. (**D**) Quantification of three biological replicates from (C). Protein levels were analyzed by densitometry on a Western blot. Data in (B) and (D) are shown as mean ± s.d. P values determined by one-sample t-test. *p<0.05, *p<0.005

Next, we aimed to determine whether PARP7 MARylation regulates p300 stability in a UPS-dependent manner in CT26 cells, given that our previous studies, along with those of others, demonstrated that PARP7 MARylation modulates the type I IFN response in these cells (*7, 19*). To achieve this, we generated CT26 cell lines that stably express doxycycline (dox)-inducible GFP-tagged WT-PARP7 or ΔKBR3-PARP7. To exclude effects from endogenous PARP7, these stable lines were established in a PARP7 KO background. Experiments were performed using the STING agonist DMXAA, which is known to promote IRF3 phosphorylation and drive subsequent transcriptional activation through IRF3–p300/CBP complexes (*42*). Like we observed in HEK 293T cells, dox-induction of WT-PARP7, but not ΔKBR3-PARP7 treated with DMXAA (10 µg/mL), led to a statistically significant reduction in p300 levels, which was prevented by treatment with proteosome inhibitor MG132 (5 μM) (Fig. 4C, D). p300 levels were unaffected by MG132 in the PARP7 KO cells. Overall, these results strongly support a model where PARP7 MARylation facilitates UPS-dependent p300 degradation.

### p300/CBP regulates auto-MARylation and PARP7 nuclear foci

The activity of a PTM enzyme can be influenced by its interaction with a substrate, including some members of the PARP family members (*43, 44*). This led us to ask if PARP7 binding to p300/CBP regulates the catalytic activity of PARP7 in HEK 293T cells. To test this idea, we initially compared the auto-MARylation activity of GFP-tagged WT- versus ΔKBR3- PARP7 in HEK 293T cells. We found that WT-PARP7 auto-MARylation was 2.5-fold lower than that of ΔKBR3-PARP7 after normalizing the auto-MAR signal to total GFP-tagged PARP7 levels (Fig. 5A, B). Since KBR3 deletion could exert effects independent of p300/CBP binding, we then tested if depletion of p300/CBP using the small molecule-based p300/CBP PROTAC, dCBP-1 (49), impacts the catalytic activity of PARP7. We found that treatment of cells with dCBP-1 (1 μM), which effectively depleted p300/CBP, resulted in a statistically significant decrease in WT-, but not ΔKBR3, auto-MARylation (Fig. 5C, D). This effect was not observed upon treatment with A485 (10 μM) (*45*), an inhibitor of the histone acetyltransferase activity of p300/CBP (Fig. 5C, D).

**Fig. 5.**
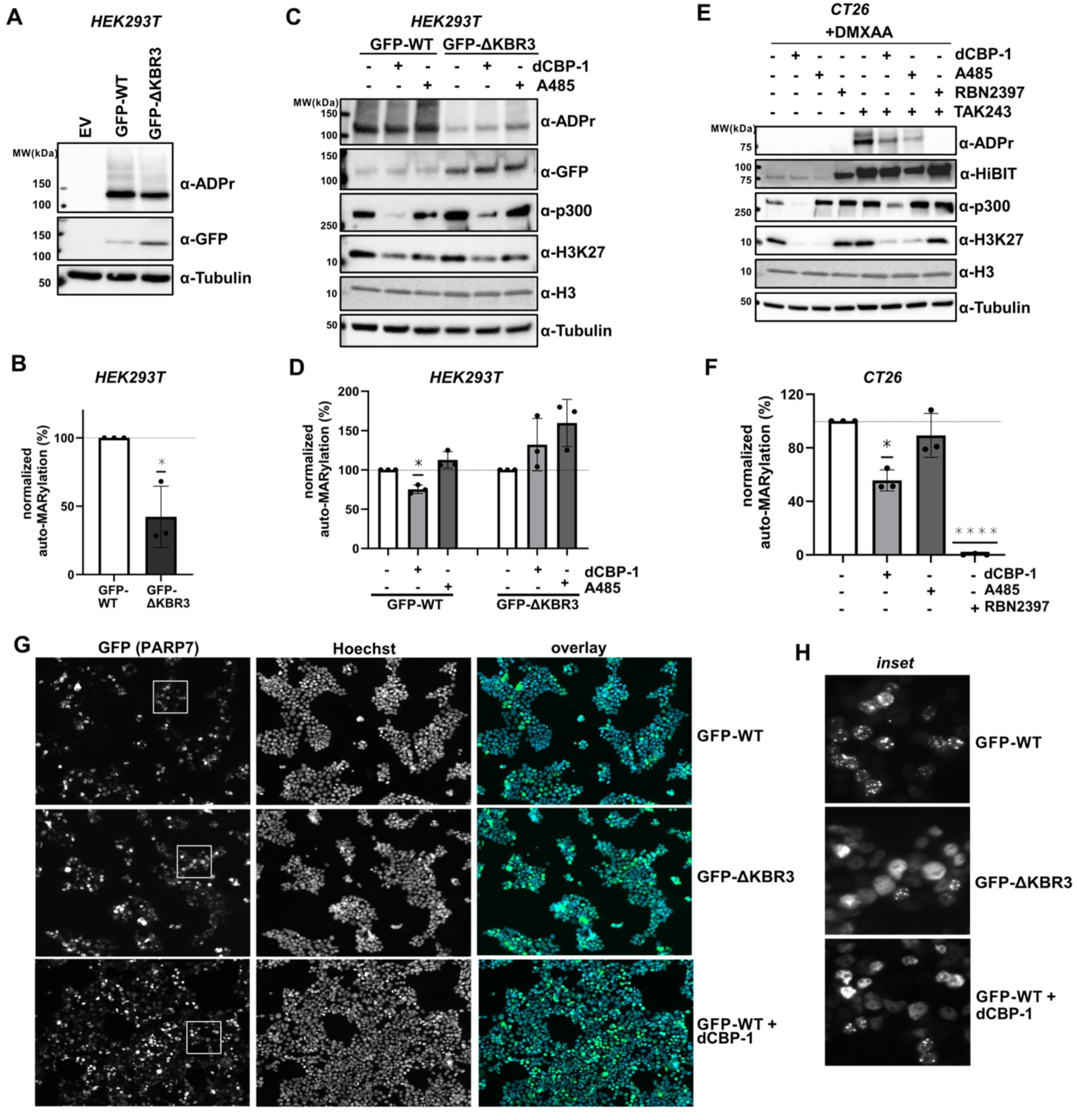
p300/CBP regulates auto-MARylation and PARP7 nuclear foci. (**A**) Western blot showing PARP7 auto-MARylation and expression in HEK 293T cells expressing GFP-PARP7-WT and -ΔKBR3. Cells were treated with 100 nM AZD5305 to avoid PARP1-mediated PARylation signals, which occur near the molecular weight of GFP-PARP7. (**B**) Quantification of the levels of ADP-ribosylation in (A). (**C**) Western Blot showing the effect of p300 inhibition/degradation on PARP7 auto-MARylation in HEK 293T cells expressing GFP-PARP7-WT and ΔKBR3 ± 10 μM A485 or 1 μM dCBP for 24 h. (**D**) Quantification of three biological replicates from (C). (**E**) Effect of p300 inhibition/degradation on PARP7 auto-MARylation in HiBiT-PARP7 KI CT26 cells ± 10 μM A485 or 1 μM dCBP. Cells were treated with 1 uM TAK243 to prevent PARP7 degradation and 100 nM AZD5305 to avoid PARP1 background. (**F**) Quantification of three biological replicates from (E). Protein and ADPr levels were analyzed by densitometry on a Western blot. (**G**) IF analysis showing how expression of PARP7-ΔKBR3 and degradation of p300/CBP impact PARP7 localization compared to untreated PARP7-WT. HEK 293T cells were transiently transfected with GFP-PARP7-WT and GFP-PARP7-ΔKBR3 and treated with DMSO or transfected with GFP-PARP7-WT and treated with 1 uM dCBP-1. (**H**) Representative images from (G).

We next asked whether dCBP-1 exerted similar effects on endogenous PARP7 auto- MARylation. For these experiments, we used our previously described knock-in (KI) CT26 cell line in which endogenous PARP7 was N-terminally tagged with an 11 amino acid peptide (HiBiT) (*12*). These KI cells allowed facile detection of endogenous PARP7 using an anti-HiBiT antibody. Endogenous PARP7 auto-MARylation could not be detected in DMXAA-treated HiBiT-PARP7 KI CT26 cells; however, treatment with the ubiquitin-like modifier activating enzyme 1 (UBA1) inhibitor, TAK243 (1 μM), which substantially increased endogenous PARP7 levels, revealed an ADPr band just above ∼75 kDa, the molecular weight of HiBiT-PARP7 (Fig. 5E). Treatment with RBN2397 blocked the TAK243-induced ADPr band, consistent with the idea that this ADPr band is likely auto-MARylated endogenous PARP7 (Fig. 5E). dCBP-1, but not A485, significantly decreased normalized (i.e., relative to total HiBiT-PARP7 levels) auto- MARylated endogenous PARP7 (Fig. E, F). These observations for endogenous PARP7 in HiBiT-PARP7 KI cells, combined with data for GFP-tagged WT- and ΔKBR3-PARP7 in HEK 293T cells, provide evidence that the catalytic activity of PARP7 is regulated by its interaction with p300/CBP.

Upon expression in multiple cell types, PARP7 localizes to foci in the nucleus (*12, 15*). Catalytically inactive PARP7 mutants or treatment with the PARP7 inhibitors RBN2397 and KMR206 disrupt foci and lead to diffuse nuclear localization (*12*), supporting the idea that PARP7 MARylation regulates its localization to nuclear foci. Given the effects of p300/CBP on PARP7 MARylation, we subsequently explored whether PARP7’s interaction with p300/CBP impacts its localization. We found that either the deletion of KBR3 or the treatment with dCBP-1 disrupted PARP7’s localization to nuclear foci (Fig. G, H). These observations corroborate our earlier results, providing further evidence that p300/CBP binding to PARP7 regulates its catalytic activity.

### PARP7 KBR3 is required for PARP7-dependent suppression of IFNβ expression

Considering the critical function of the KBR3 α-helix of PARP7 in mediating PARP7’s interaction with and MARylation of p300/CBP—and the established role of p300/CBP as a key co-factor for IRF3—we next sought to examine the functional impact of deleting KBR3 on PARP7-mediated suppression of IRF3-driven transcription of *IFNβ*. As an initial approach, we used a previously described luciferase-based IFNβ reporter assay in HEK 293T cells to assess the effects of PARP7 on IRF3-driven transcription. In this assay, PARP7 (and PARP7 variants) are co-expressed with a hemagglutinin (HA)-tagged, constitutively active variant of IRF3, HA-IRF3- 5D (*46*), enabling the assessment of the suppressive effects of PARP7 on IRF3-driven transcription, independent of upstream activation. Consistent with recent studies, we found that co-expression of GFP-WT-PARP7 with HA-IRF3-5D completely abolished IFNβ-dependent luciferase activity (Fig. 6A). By comparison, GFP-ΔKBR3-PARP7 did not completely suppress IFNβ-dependent luciferase activity (Fig. 6A). Despite the striking effect of KBR3 deletion on p300/CBP MARylation and binding to PARP7, the overall impact on PARP7-mediated suppression of IFNβ-dependent luciferase activity was less pronounced than expected. We therefore evaluated the effects of PARP7 inhibitors, RBN2397 or KMR206, on PARP7-mediated suppression of IFNβ-dependent luciferase activity in cells expressing WT- and ΔKBR3-PARP7. Treatment of WT-PARP7 expressing cells with saturating doses of either RBN2397 (300 nM) or KMR206 (300 nM) partially reversed the suppressive effects of WT-PARP7 on IFNβ-dependent luciferase activity to levels like those observed in ΔKBR3-PARP7 expressing cells (Fig. 6B). By contrast, neither PARP7 inhibitor impacted the suppressive effects of ΔKBR3-PARP7 (Fig. 6B). Together, these results show that deletion of KBR3 relieves the inhibitor-sensitive PARP7- mediated suppression of IFNβ-dependent luciferase activity. Thus, we propose that the catalytic activity-dependent suppressive effects of PARP7 on IRF3-driven transcription depend on PARP7’s interaction with p300/IRF3. Conversely, the catalytic activity-independent effects of PARP7 on IRF3-driven transcription appear independent of PARP7 binding to p300/CBP.

**Fig. 6.**
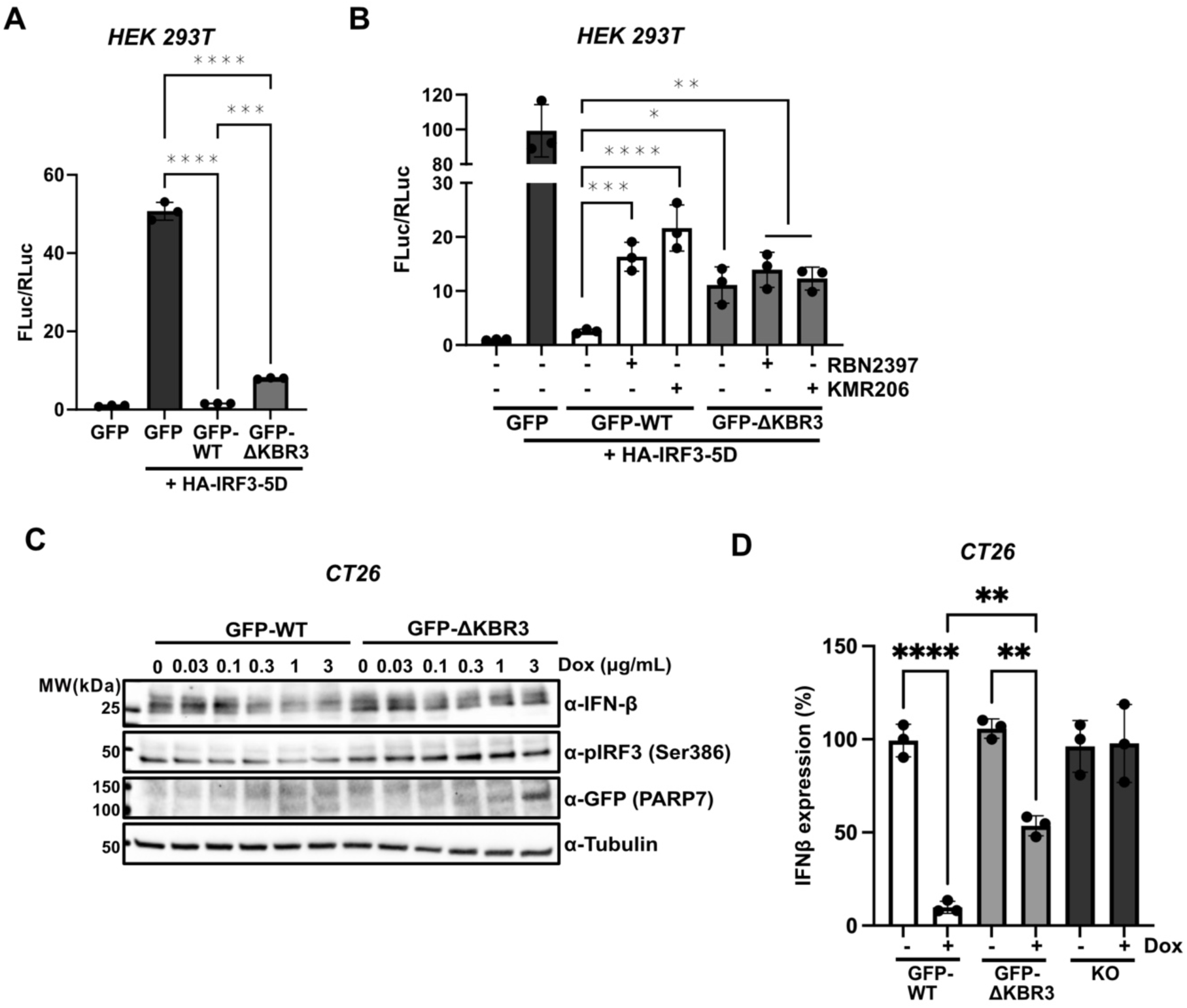
PARP7 KBR3 is required for PARP7-dependent suppression of IFNβ expression. (**A**) Quantification of luciferase activity in HEK 293T cells transfected with *IFNB1*-FLuc, TK- RLuc, HA-IRF3-5D alone or in combination with GFP-PARP7-WT and GFP-PARP7-ΔKBR3 for 24 h. (**B**) Quantification of luciferase activity in HEK 293T cells transfected with *IFNB1*-FLuc, TK- RLuc, HA-IRF3-5D alone or in combination with GFP-PARP7-WT or GFP-PARP7-ΔKBR3 and treated with either DMSO, 300 nM RBN2397, or 300 nM KMR206 for 24 h. (**C**) Western blot of dox-inducible CT26 cells expressing PARP7-WT or ΔKBR3 incubated with increasing concentrations of Dox for 24 h and stimulated with 10 ug/mL DMXAA for 6 h before harvest. (**D**) ELISA of mouse IFNβ in the supernatant of PARP7-KO CT26 cells, and CT26 expressing WT- and ΔKBR3- PARP7 treated with 5 μg/mL Dox for 24 h, and 10 ug/mL DMXAA for 6 h. Data in (A), (B), and (D) is shown as mean ± s.d. *P* values were determined by one-way ANOVA. \**p*<0.05, \*\**p*<0.01, \*\*\**p*<0.001, \*\*\*\**p*<0.0001.

We turned our attention to examining the effects of KBR3 deletion on endogenous IFN-β expression levels. We measured the levels of IFNβ by western blot in dox-inducible WT-PARP7 or ΔKBR3-PARP7 CT26 cells treated with DMXAA and increasing concentrations of dox. We observed a dose-dependent decrease in IFNβ levels in WT-PARP7, but not PARP7-ΔKBR3 cells, confirming earlier findings about the suppressive role of WT-PARP7 (Fig. 6C). Notably, phosphorylated IRF3 (phospho-Ser386) levels declined in parallel with IFNβ levels with increasing dox concentrations in WT-PARP7 cells (Fig. 6C). This increase in phospho-Ser386 IRF3 was not observed in ΔKBR3-PARP7 cells (Fig. 6C). We also measured the levels of secreted IFNβ in dox-inducible WT-PARP7 or ΔKBR3-PARP7, as well as PARP7 KO CT26 cells treated with DMXAA, using an ELISA. Treatment of WT-PARP7 cells with dox reduced IFNβ levels ∼13-fold (Fig. 6D). By contrast, dox treatment only modestly reduced (∼2-fold) IFNβ levels in ΔKBR3-PARP7 (Fig. 6D). Importantly, dox treatment did not impact IFNβ levels in PARP7 KO CT26 cells (Fig. 6D). Collectively, these results indicate that PARP7’s association with p300/CBP is essential for achieving the full reduction in IFNβ expression, aligning with a role for PARP7 in the transcriptional repression of IRF3 in a p300/CBP-dependent manner.

### p300/CBP-dependent interaction underlies PARP7 inhibitor–mediated enhancement of IFNβ expression

Typically, the phenotypic effects of active site-directed, small-molecule inhibitors of enzymes in cells arise from inhibition of catalytic function. However, in some instances, these inhibitors induce effects beyond catalytic inhibition, resulting in more pronounced or distinct phenotypes than those observed in KO models (*47, 48*). Our previous work comparing the effects of RBN2397 and KMR206 on IFNβ signaling in CT26 cells prompted us to hypothesize that, in specific cellular contexts, PARP7 inhibitors may exhibit a “dominant-positive” effect (*12*).

However, we did not directly compare the effects of PARP7 inhibitor treatment and PARP7 knockout on IFNβ signaling. Moreover, we wondered whether PARP7’s interaction with p300/CBP could be functionally relevant here. To explore this further, we evaluated the effect of PARP7 inhibition with RBN2397 on IFNβ levels in dox-inducible WT-PARP7, ΔKBR3-PARP7, and PARP7 KO CT26 cells by western blot. In cells treated with DMXAA, RBN2397 dramatically increased IFNβ levels in WT-PARP7 compared to PARP7 KO cells, whereas RBN2397 only had a marginal effect in dox-treated ΔKBR3-PARP7 cells (Fig. 7A). To ensure that the dominant-positive mechanism of PARP7 inhibitors is not a consequence of GFP-tagged PARP7 overexpression in our dox-inducible CT26 cells or specific to a given PARP7 inhibitor, we evaluated the effects of RBN2397 and KMR206 on IFNβ levels in the parent and PARP7 KO CT26 cells. Consistent with results in dox-inducible GFP-WT-PARP7 cells, IFNβ levels were significantly upregulated by RBN2397 or KMR206 treatment in DMXAA-stimulated native CT26 cells, exceeding the levels observed in PARP7 KO cells (Fig. S3). Together, these results support the notion that the PARP7 inhibitor–PARP7 complex increases IFNβ expression beyond that observed in PARP7 KO cells via a dominant positive effect.

**Fig. 7.**
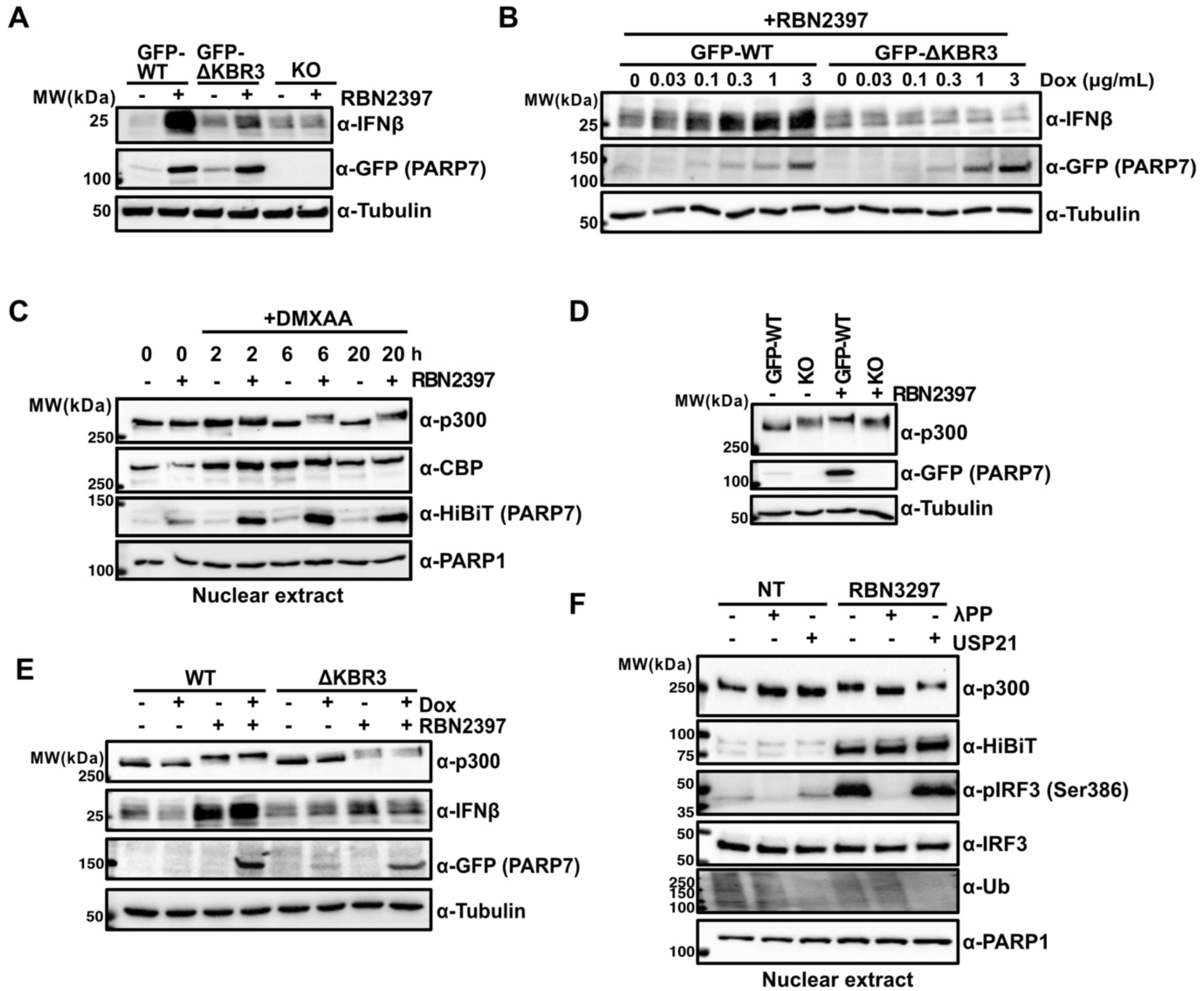
PARP7 inhibitors enhance IFNβ expression in a p300/CBP-dependent manner via a dominant positive effect and stabilize hyperphosphorylated p300. (**A**) Western blot of the levels of IFNβ in dox-inducible CT26 cells expressing PARP7-WT or ΔKBR3 ± 300 nM RBN2397 for 24 h and stimulated with 10 ug/mL DMXAA for 6 h before harvest. (**B**) Western blot of CT26 cells expressing WT and ΔKBR3 PARP7 incubated with increasing concentrations of Dox for 24 h + 300 nM RBN2397 for 24 h and stimulated with 10 ug/mL DMXAA for 6 h before harvest. (**C**) Western blot of HiBiT-PARP7 KI CT26 cells ± 300 nM RBN2397 treated with 10 ug/mL DMXAA for 0, 2, 6, and 20 h before harvest. Cells were fractionated, and nuclear extracts were loaded in 6% SDS-PAGE to achieve optimal band separation. (**D**) Western blot of CT26 cells expressing WT PARP7 or PARP7-KO CT26 cells treated with 3 ug/mL Dox for 24 h ± 300 nM RBN2397. Cells were stimulated with 10 ug/mL DMXAA for 6 h with before harvest. Cells lysates were loaded in 6% SDS-PAGE for analysis. (**E**) Western blot of IFNβ in dox-inducible CT26 cells expressing PARP7-WT or ΔKBR3 treated with 3 ug/mL Dox for 24 h ± 300 nM RBN2397. Cells were stimulated with 10 ug/mL DMXAA for 6 h. Cells lysates were loaded in 6% SDS-PAGE for analysis. (**F**) Western blot of HiBiT-PARP7 KI ± 300 nM RBN2397 and stimulated with 10 ug/mL DMXAA for 6 h before harvest. Nuclei of each condition were fractionated, separated in three tubes, and individually treated with either buffer, λPP, or USP21 for 30 min at 37 °C. Preparations were loaded in 6% SDS-PAGE for analysis.

In the dominant-positive model, the extent of the inhibitor-driven phenotype can be linked to the intracellular abundance and stability of the protein–inhibitor complex (*48, 49*). Based on previous inhibitor studies, we proposed that the magnitude of PARP7 inhibitor-induced IFNβ expression correlates with the levels of the PARP7 inhibitor–PARP7 complex (*12*). To further test this hypothesis—and the relevance of PARP7’s interaction with p300/CBP—we evaluated IFNβ expression in DMXAA- and RBN2397-treated dox-inducible WT-PARP7 or ΔKBR3-PARP7 CT26 cells, titrating dox doses to modulate the levels of PARP7 expression. In the presence of DMXAA and RBN2397, we found that increasing dox concentrations dose-dependently increased IFNβ levels (Fig. 7B). The rise in IFNβ levels was well-correlated with WT-PARP7 expression levels. These effects were not observed in PARP7-ΔKBR3 cells (Fig. 7B). Collectively, these findings reinforce the dominant-positive mechanism for the impact of PARP7 inhibitors on IFNβ expression levels and highlight the importance of the PARP7–p300/CBP interaction in mediating this response.

### RBN2397 stabilizes a hyperphosphorylated form of p300 in the context of STING activation

Curiously, we observed that the band for p300 shifted to a higher molecular weight form in a time-dependent manner upon treatment with RBN2397 and DMXAA in HiBiT-PARP7 KI CT26 cells (Fig. 7C). To determine whether this effect was due to RBN2397-mediated inhibition of PARP7’s enzymatic activity or a predominant positive mechanism, we compared the p300 mobility shift induced by DMXAA treatment in dox-inducible WT-PARP7 cells and PARP7 KO cells with or without RBN2397. In the absence of RBN2397, the band for p300 shifted to a higher molecular weight and appeared more diffuse in PARP7 KO cells compared to WT-PARP7 cells (Fig. 7D). When treated with RBN2397, p300 migrated as a more distinct high molecular weight band in WT-PARP7 cells than in PARP7 KO cells (Fig. 7D). We then compared the p300 molecular weight band shift in WT- versus ΔKBR3-PARP7 cells in the presence or absence of RBN2397. Following RBN2397 treatment, p300 exhibited a sharper high molecular weight band in WT cells compared to ΔKBR3-PARP7 cells (Fig. 7E). Collectively, these results suggest that PARP7 catalytic activity suppresses the formation of a higher molecular weight form of p300 upon STING activation with DMXAA, while the RBN2397–PARP7 complex promotes its stabilization via direct interaction with p300.

The shift to a higher molecular weight form of p300 suggests that p300 undergoes post-translational modification (PTM) upon STING activation with DMXAA. PTMs that would induce this dramatic shift include phosphorylation and ubiquitylation (*50–52*). To determine if either of these PTMs accounts for the DMXAA-mediated p300 molecular weight shift, we treated nuclear lysates derived from HiBiT-PARP7 KI cells with the following enzymes: 1. lambda phosphatase (λPP), a pan-Ser/Thr/Tyr phosphatase for assessing phosphorylation, or 2. the promiscuous deubiquitinase, USP21 (*53*). We found that λPP, and not USP21, abolished the DMXAA-mediated p300 molecular weight shift in RBN2397-treated cells (Fig. 7F). These results demonstrate that phosphorylation underlies the molecular weight shift of p300 in STING-activated and RBN2397-treated CT26 cells.

## Discussion

Although PARP7 has been identified as a key suppressor of IFNβ signaling in specific tumor cells, a detailed understanding of the underlying mechanisms has remained elusive, especially regarding the role of PARP7-mediated MARylation. Herein, we developed an optimized ASCG approach for identifying low-abundant, direct substrates of PARP7. The nuclear substrates we identified include many proteins associated with transcriptional regulation and chromatin remodeling, supporting PARP7’s known function in ligand-dependent transcriptional control (*11, 15–18, 31, 54*). Although we identified transcription factors, such as AHR and AR, as substrates of PARP7, confirming previous studies that have shown these to be substrates of PARP7 (*11, 55*), the predominantly enriched nuclear substrates were transcriptional co-regulators, including co-activators and co-repressors. In this study, we focused on the co-activators p300 and CBP because (i) they are top-ranked PARP7 interactors (*19, 32*), and (ii) they are key regulators of IRF3-driven *IFNβ* transcription (*30, 56*). Guided by AlphaFold modeling, we demonstrate that the formation of the PARP7–p300/CBP complex in cells is mediated by a novel interaction between an α-helix (which we call KBR3) in PARP7’s N-terminus and the KIX domain of p300/CBP. Our findings in two cellular contexts using both WT-PARP7 and the ΔKBR3 variant support a model in which the PARP7–p300/CBP complex regulates type I interferon signaling via a combination of loss-of-function and inhibitor-mediated dominant positive mechanisms (Fig. 8). In the loss-of-function mechanism, PARP7-mediated MARylation of p300/CBP targets them for proteasomal degradation, leading to the loss of IRF3-dependent transcription of *IFNβ* (Fig. 8A).

**Fig. 8.**
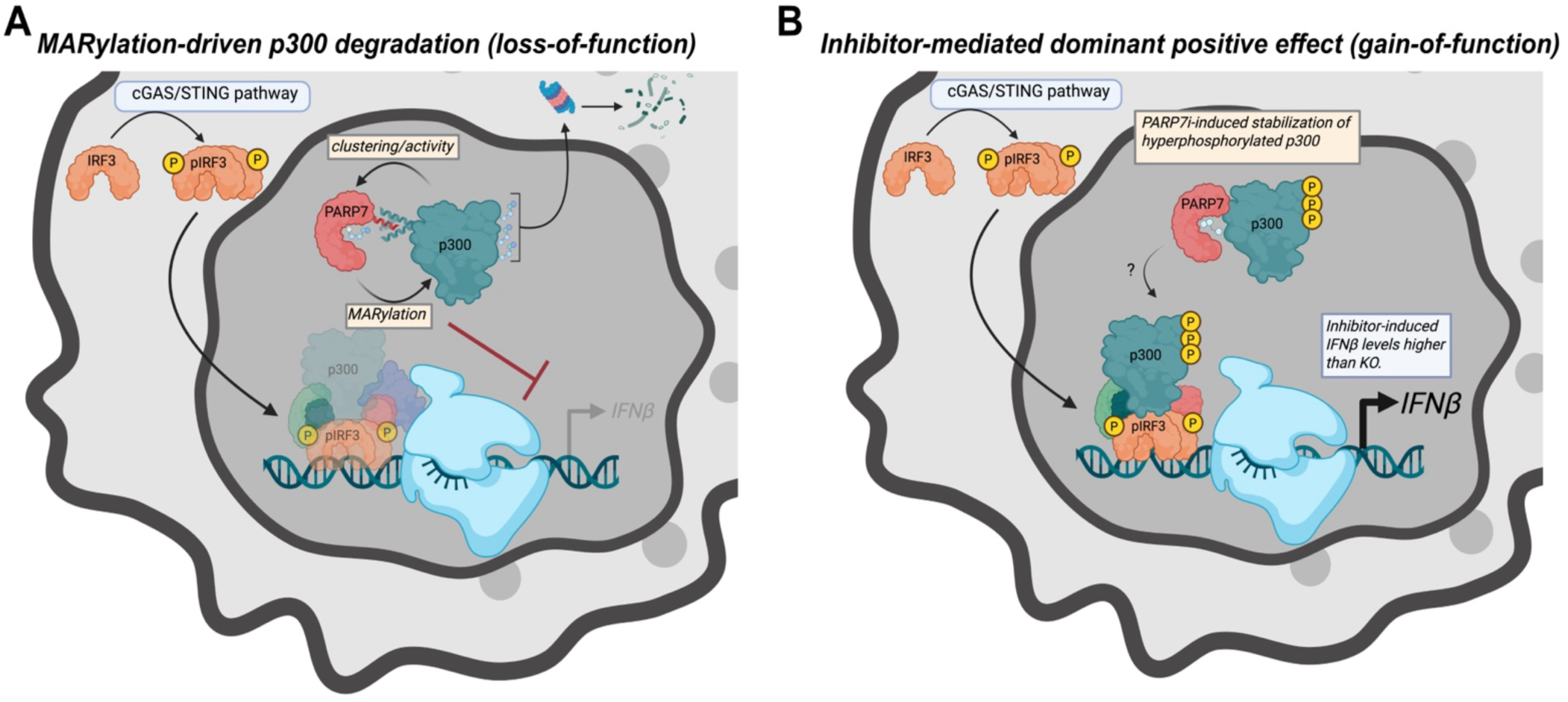
Both loss- and gain-of-function mechanisms account for the regulation of IFNβ expression by PARP7 inhibitors. In both mechanisms, p300 (and likely CBP) play a critical role through interactions with PARP7.

Recent studies suggested that PARP7 interacts with IRF3 and disrupts the interaction between p300/CBP and IRF3 (*8*) Although not mutually exclusive with our model, this loss-of-function mechanism could be explained by the decreased phospho-IRF3 levels in the nucleus of PARP7 KO cells compared to WT cells (*8*) an effect we observe when we compare DMXAA-stimulated WT- versus ΔKBR3-PARP7 CT26 cells. This suggests that PARP7 may regulate nuclear phospho-IRF3 indirectly through its interaction with p300/CBP.

An intriguing, unexpected aspect of this mechanism is that p300/CBP regulates PARP7 auto-MARylation and localization in nuclear foci (Fig. 8A). These effects depend on physical interactions with p300/CBP, rather than their catalytic activity. This mode of substrate-driven regulation is similar to that observed for the PARP1 substrate HPF1; HPF1 binding to PARP1 increases PARP1’s catalytic activity and alters its amino acid targeting specificity on substrates (*43, 57*).

Finally, our results support the notion that PARP7 inhibitors increase IFNβ levels beyond those observed with genetic deletion of PARP7 via a dominant positive effect. This dominant positive effect is dependent on PARP7’s interaction with p300/CBP (Fig. 8B). Prolonged treatment with PARP7 inhibitors leads to the stabilization of a hyperphosphorylated state of p300 (Fig. 8B). Overall, our study provides new insight into the mechanism by which PAPR7 acts as a potent suppressor of IFNβ signaling in tumor cells.

A key finding in our study is the identification of three helices, KBR1-3, located in the N-terminus of PARP7, which are necessary for PARP7 binding to p300. Among the three helices, KBR3 exhibited the most extensive interaction and the lowest predicted alignment error, providing confidence in the predicted interface between PARP7 and the c-Myb site of the KIX domain of p300. Moreover, KBR3 contained the conserved KIX binding motif ΦXXΦΦ. Given that other transcription factors, such as c-Myb and E-protein, bind to the same site, it will be interesting to determine whether PARP7 directly competes with these transcription factors at the c-Myb site to regulate their transcriptional activity. Another possibility is that binding a transcription factor to the MLL site (e.g., MLL, CREB) enhances binding of KBR3, promoting the formation of a ternary complex between PARP7–p300–transcription factor. Indeed, MLL binding to the KIX domain cooperatively enhances the binding of c-Myb to the KIX domain (*58*). Finally, it will be interesting to determine if KBR1 and/or KBR2 contribute to PARP7 binding to p300.

Beyond KBR3, other domains of PAPR7 are also necessary for regulating IRF3-driven transcription of IFNβ, as shown using an IFNβ reporter assay in HEK 293T cells (*8*) Indeed, deletion of the ZnF or the second WWE (WWE2) domain reverses PARP7-dependent inhibition of IFNβ-induced transcription, despite not affecting IRF3 binding (*8*) While the exact function and binding partners of these domains are not yet known, they appear to play a role in the nuclear localization of PARP7 (*59*). Therefore, their impaired ability to suppress IFNβ-induced transcription may stem from a decreased localization to the nucleus and, consequently, a reduced interaction with p300/CBP.

Our finding that PARP7’s catalytic activity regulates the stability of p300 in a UPS-dependent manner aligns with previous studies demonstrating similar regulation of other substrates, including various transcription factors (*15, 16, 54*). Taken together, these results suggest the intriguing idea that PARP7-mediated MARylation directs ubiquitin-dependent degradation of PARP7 substrates. How might PARP7-mediated MARylation regulate protein stability via the ubiquitin-proteasome system? Classically, ubiquitylation involves the covalent attachment of ubiquitin’s (Ub) carboxyl terminus to lysine (Lys) residues on substrate proteins, forming an isopeptide bond (*60*). The initial Ub mark can be further modified to form a Ub polymer (polyUb). The diversity of ubiquitylation signaling stems from the individual or multiple Lys residues of Ub used to extend the polymer. The primary polyUb linkage responsible for protein degradation is K48-linked chains (*61*). However, other polyUb chains, like K11-linked bonds (*62*) and K11/48-branched chains (*63*), can also signal for degradation. Ubiquitylation occurs through a cascade involving E1 Ub-activating, E2 Ub-conjugating, and E3 Ub-ligating enzymes. Our study using proximity biotinylation proteomics (BioID) (*19*), along with an independent co-immunoprecipitation study (*15*), identified several PARP7-interacting E3 ligases, including HUWE1, UBR5, RNF114, and DELTEX (DTX) DTX2. In the case of RNF114 and DTX2, they have been shown to bind to MARylated peptides and substrates (*64–66*). In another study, HUWE1 was shown to interact with PARP7 in a MARylation-dependent manner and was suggested to regulate the stability of other PARP7-interacting proteins (*15*). Future work will determine whether any of these E3 ligases regulate p300 stability in a PARP7 catalytic activity-dependent manner, and if so, whether MAR conjugation to the substrate serves as a degradation mark.

Recent studies from several groups show that p300/CBP can form biomolecular condensates in the nucleus (*67–69*). These condensates can sequester nucleosome components, and a fraction of these condensates co-localize with repressive histone marks, namely H3K27Me3, potentially acting as a storage pool of p300 at silenced promoters (*67*). p300 can also drive the formation of co-condensates with several transcription factors (*68*) and oncogenic fusion proteins (e.g., BRD4–NUT) (*70*). Our observation that PARP7 nuclear foci formation depends on its interaction with and the presence of p300 is consistent with these findings. Given that PARP7’s catalytic activity is necessary for its nuclear foci formation, and our findings that disrupting the PARP7–p300 interaction or loss of p300 diminishes PARP7 auto-MARylation, we propose that p300 promotes co-condensate formation with PARP7 by either enhancing PARP7 catalytic activity or stabilizing auto-MARylated PARP7. Intriguingly, a recent study showed that PARP14 forms cytoplasmic co-condensates with its putative substrate p62 (also known as sequestosome 1), which depends on p62 and the catalytic activity of PARP14 (*71*). Thus, MARylation-dependent co-condensation of PARPs with their substrates may constitute a general mechanism by which cells achieve spatial and temporal regulation of PARP activity.

In previous work, we demonstrated that the magnitude of PARP7 inhibitor-induced IFNβ expression in CT26 cells correlated with the effects of the inhibitor on PARP7 protein levels (*12*). This observation, together with recent studies demonstrating that loss of PARP7 abolishes the impact of RBN2397 on cell viability (*7, 14*), led us to hypothesize that, beyond a loss-of-function mechanism, PARP7 inhibitors act through a dominant positive effect—where the PARP7 inhibitor–PARP7 complex drives the cellular phenotype of PARP7 inhibitors (*72*). By showing that RBN2397 enhances IFNβ levels beyond what would be expected from PARP7 loss alone, our current study substantiates the dominant positive model underlying the cellular effects of PARP7 inhibitors. Additionally, we demonstrate that PARP7 inhibitor-induced IFNβ expression requires PARP7’s interaction with p300. Collectively, our findings offer more detailed mechanistic insight into how PARP7 inhibitors exert a dominant positive effect on IFNβ expression. Future studies will focus on elucidating the molecular features of the PARP7 inhibitor–PARP7–p300 complex and how this complex drives IFNβ expression. The dominant positive effect of PARP7 inhibitors on IFNβ expression is reminiscent of the impact of positive allosteric modulators of PARP1 binding to DNA and subsequent cell death.

Another notable finding in our study was the persistence of a slower-migrating form of p300 in the presence of the STING agonist DMXAA and PARP7 inhibitors. This form of p300 was present in PARP7 KO, though the mobility shift was less pronounced and weaker in magnitude. Similar results were observed in CT26 cells expressing ΔKBR3-PARP7 compared to WT-PARP7. Treatment of lysates with λPP revealed that the slower-migrating form of p300 is indeed hyperphosphorylated (due to the dramatic molecular weight shift) p300. Based on these results, we propose that PARP7 catalytic activity either suppresses the activity of a p300 phosphatase or activates a p300 kinase, and that PARP7 inhibitors stabilize hyperphosphorylated p300 via a dominant positive effect. While no phosphatase has been shown to interact with p300, several kinases have been reported to phosphorylate p300, including CDK1, ERK1/2, PKC, ATM, and mTOR (*73–77*). Previous studies have shown that p300 phosphorylation regulates its stability, transcriptional activity, and localization depending on cellular context (*73, 74, 76, 77*). Intriguingly, all but two (ERK1/2) of these kinases and the regulatory subunit of protein phosphatase 1 (PP1), PP1R10, were identified as substrates of PARP7 using our ASCG approach (Table S1). Follow-up studies will investigate potential roles for these kinases and PP1R10 in regulating p300 phosphorylation under PARP7 inhibitor treatment. We will also determine whether p300 phosphorylation is necessary for PARP7-inhibitor induced IFNβ expression, and if so, delineate its function.

While our findings provide new insight into the mechanism by which PARP7 regulates IFNβ expression, several limitations should be acknowledged. First, although our ASCG approach using HEK 293T cell lysates identified functionally relevant substrates such as p300/CBP, it is possible that we did not capture additional substrates that are specifically expressed in CT26 cells. Moreover, MARylation of some PARP7 substrates likely requires activation of cGAS/STING pathway. Further optimization of the expression of our analog-sensitive PARP7 variant in CT26 cells is necessary before applying the ASCG approach in these cells, especially under cGAS/STING activation. Second, our studies indicate a dominant-positive effect of PARP7 inhibitors on IFNβ expression in CT26 cells; whether this effect extends to other tumor types remains to be determined. Finally, how PARP7 inhibitors drive IFNβ expression through a dominant positive effect remains to be explored. Does the PARP7 inhibitor–PARP7–p300 complex localize to the IFNβ promoter? Does this complex stabilize IRF3 at the promoter or lead to the formation of a more transcriptionally active complex? These and other questions will be addressed in due course.

In summary, we demonstrated that PARP7 negatively regulates IFNβ expression through its interaction with and MARylation of the transcriptional co-activator p300, leading to p300 degradation. Additionally, we show that PARP7 inhibitors exhibit a dominant positive effect, driving high levels of IFNβ expression. These findings not only provide a deeper understanding of the mechanism by which PARP7 MARylation regulates gene transcription but also hold important implications for the therapeutic advancement of PARP7 inhibitors.

## Methods

### Cell culture

HEK 293T were maintained in DMEM supplemented with 10% fetal bovine serum (FBS) and 1X Glutamax. Transfection with plasmids was performed using JetOPTIMUS transfection reagent according to the manufacturer’s protocol. CT26 cells were maintained in RPMI media and added with 10% FBS. For transient transfection cells were seeded 16 h before transfection (70000/cm^2^) in a 10 cm dish. Then, cells were transfected using 6 μg of expression vector using jetOptimus (Polyplus) according to the transfection manual using a 1:1.5 ratio of DNA/transfection reagent. The medium was exchanged and replaced with fresh medium after 4-6 h. Gene expression was allowed to proceed for 24 h. Then, the cell dishes were washed with 1X PBS, the PBS was removed, and the dish was stored at –80 °C until further use. For the gene expression of GFP-wt- PARP7, GFP-IG-PARP7, and empty vector (EV)-GFP, pEGFPC1vectors were used as previously published (19).

### ASCG labeling in HEK 293T lysates using 5-Bn-DTB-NAD⁺ and enrichment with NeutrAvidin

Lysis buffer (25 mM HEPES, 50 mM NaCl, 5 mM MgCl_2_, 1% (v/v) NP40 1, 1 mM TCEP, 1X cOmplete EDTA-free protease inhibitor (Roche), 200 μL) was added to each HEK 293T dish transfected with GFP-wt-PARP7, GFP-IG-PARP7, or EV-GFP and the dish was incubated on ice for 15 min. Subsequently, the cell lysate was transferred to an Eppendorf tube, sonicated for 3 × 5 s and the cell debris was cleared by centrifugation (4°C, 14,000 x g, 15 min). The cell lysates were transferred to a new Eppendorf tube, and the protein concentration was determined via BCA (Pierce™). The protein concentration was adjusted to 2 mg/ml with lysis buffer, 5-Bn- DTB-NAD⁺ (100 μM) was added (total volume: 500 μL per condition) and the reaction mixture was incubated for 2 h (30 °C, 400 rpm). The reaction was then allowed to cool to room temperature for 5 min and transferred to a Falcon containing MeOH (-20 °C, 4 mL), CHCl_3_ (1 mL), and Milli-Q water (2.5 mL). The Eppendorf tube was washed with 1 mL ice-cold Milli-Q water, the water was added to the falcon and the mixture was centrifuged (4°C, 5000 x g, 10 min). After centrifugation, a protein disc was visible between the CHCl_3_ and MeOH/water layers. The solvent was carefully removed without disturbing the protein disc, and the disc was resuspended in cold MeOH (1 mL). The suspension was transferred to a new Eppendorf tube, sonicated or vortexed briefly and the proteins were pelleted by centrifugation (4°C, 5000 g, 10 min). MeOH was removed, and cold MeOH (1 mL) was added. The proteins were then resuspended and centrifuged as before. MeOH was removed, and the protein pellet was air-dried for 15 minutes.

Then, 6 M urea in 1X PBS (500 μL) and 10% (w/v) SDS (25 μL) was added. The protein pellet was sonicated and heated at 37°C for 30 min to redissolve the proteins and 30 μL was saved as input. The redissolved proteins were then added to a low-binding tube containing 100 μL of pre- equilibrated Pierce™ High Capacity NeutrAvidin™ Agarose (Thermo Scientific™) and the mixture was incubated rotating overnight at 4 °C. The next day, the tubes were centrifuged (90 sec, 489 g) and the mixture was transferred to a Morbicol “F” column (MoBi Tec) with a 35 μm filter inserted for washing. After centrifugation (90 sec, 489 g) 30 μL were saved as unbound. Subsequently, the beads were washed with 0.2% (w/v) SDS in 1X PBS (500 μL, 5×) and 1X PBS (5 ×). Then, 1X PBS (500 μL) was added and the beads were transferred to a fresh low-binding tube and centrifuged (90 sec, 489 g). The supernatant was removed, elution buffer (150 μL, 4% (w/v) SDS, 5 mM biotin in Milli-Q water) was added and the proteins were eluted by incubation at 95 °C (5 min, 400 rpm). The samples were centrifuged (3 min, 4000 g) and the supernatant was transferred to a new low-binding tube. 10 μL of the elution was saved as control and the residual elution was stored at −20 C until further use.

6X loading dye (225 mM Tris-HCl, pH 6.8, 50% (v/v) glycerol, 5% (w/v) SDS, 0.05% (w/v) bromophenol blue, 12.5% (v/v) β-mercaptoethanol) was added to the input, unbound, and elution controls. Controls were denatured at 95°C for 5 minutes and stored at −20°C until further analysis. The experiment was performed in triplicate.

### Protein digest

Eluted proteins were digested using S-Trap™ sample processing technology. All used buffers were prepared on the same day of the experiment. The following stock solutions were used: 1M TEAB Thermo Scientific™), MeOH (LC-MS grade, VWR), and formic acid (Optima™ LC-MS grade, Fisher Scientific™).

Samples were dried with a SpeedVac (Thermo Scientific SDP121P) for 2 h at room temperature. Subsequently, 1X protein solubilization buffer (150 μL, 5% (w/v) SDS, 50 mM TEAB) was added and the samples were vortexed for 30 sec until complete solubilization. The samples were centrifuged (18,300 x g, 3 min), DTT (500 mM, 6.8 μL) was added to each sample, and the samples were incubated at 95°C for 10 min. After cooling the samples to room temperature (10 min), iodoacetamide (500 mM, 13.6 μL) was added and the samples were incubated at room temperature in the dark for 30 min. Aqueous phosphoric acid (12%, 17 μL) was added to each tube, the samples were mixed, and S-trap binding buffer (1124 μL, 90% (v/v) MeOH, 10% (v/v) 1 M TEAB) was added. After mixing, the samples were transferred to the S- Trap™ micro columns (Pofiti). It should be noted that the samples were loaded in multiple steps, with only 164 μL being added each time. Then, the S-Trap microcolumns were centrifuged (4000 x g, 3 min), and the captured proteins were washed by adding S-Trap binding buffer (150 μL), followed by a second centrifugation (4000 x g, 3 min) of the microcolumns. The washing step was repeated five times, and the columns were placed in a fresh low-binding tube. Digesting buffer (40 μL, 50 mM TEAB, trypsin (Promega) 80 ng/μL) was added to each column, the columns were loosely sealed, and the samples were incubated overnight at 37°C. The next day, the peptides were eluted with 50 mM TEAB (40 μL), aqueous 0.2% (v/v) formic acid (40 μL), and 0.2% (v/v) formic acid in ACN/Milli-Q water (50/50, 35 μL). The micro columns were centrifuged (4000 x g, 4 min) after each elution step. The elution fractions were pooled and dried using a SpeedVac at room temperature for 1.5 h. 100 μL of MeOH was added, and the samples were vortexed and dried again with a SpeedVac at room temperature. Samples were stored at - 80°C before further usage.

### LC-MS/MS measurements and data analysis

LC-MS/MS measurements were performed by the proteomic center of the University of Konstanz. After proteolytic digestion samples were desalted using Pierce^TM^ C18 Spin Tips (Thermo Scientific) and eluted with 0,1% formic acid in ACN/Milli-Q water (80/20, 2× 20 μL). To evaporate ACN, samples were incubated for 52 min at 42 °C. Subsequently, samples were diluted with 17 μL of 0.1% formic acid in Milli-Q water and were analyzed on a QExactive HF mass spectrometer (Thermo Fisher Scientific) interfaced with an Easy-nLC 1200 nanoflow liquid chromatography system (Thermo Fisher Scientific). Samples were loaded onto the analytical column (75 μm × 50 cm C18 Acclaim PepMap, 100Å pore size) and peptides were resolved at a flow rate of 150 nL/min using a linear gradient of 6−48% solvent B (0.1% formic acid in 80% acetonitrile) over 108 min. Samples were measured in technical duplicates in data-independent acquisition. MS1 scans were acquired in a 300−1650 m/z range at a mass resolution of 120000 (at 200m/z). The whole mass range was divided in 24 variable mass windows for isolated ion fragmentation (m/z between 27 and 470, see attached excel sheet) with a MS/MS resolution of 30000. Ions were fragmented using higher-energy collision dissociation (HCD) set to 28%, AGC target: 1e6, injection time: auto, default charge: 2.

Raw files were analyzed using Spectronaut® 18 (Biognosys) (*84*) with a library-free directDIA+ deep workflow mode. The minimum peptide length was set to 7 and for identification the following values were used; Precursor Q value cutoff: 0.01, Precursor PEP cutoff:0.2 ProteinQvalue cutoff(Experiment): 0.01, ProteinQvalue cutoff(Run):0.05, Protein PEP Cutoff 0.75. As protein database, the FASTA file swiss prot UP000005640 was used. Contaminants were identified with the contaminant FASTA by Frankenfield et al. (*85*). Protein LFQ method was set to MaxLFQ and samples were normalized using the implemented Cross-Run Normalization (Filter Type: None, Strategy: Automatic).

Statistical analysis of the raw LFQ data was performed with Perseus software (1.6.15.0) (*86*). LFQ intensities were log2 transformed and filtered to be detected in at least 4 out of 6 replicates, and missing values were imputed from a normal distribution (width = 0.3 and shift = 1.8 for total matrix). Technical replicates were averaged (mean), filtered for contaminants and significantly enriched proteins were identified by ANOVA (S0 = 0.2, FDR = 0.001) followed by hierarchical clustering of significant different means (Tukey’s honestly significant difference test, FDR = 0.05) according to their Euclidean distance.

### ASCG labeling in HEK 293T intact nuclei using 5-Bn-DTB-NAD⁺ and enrichment with NeutrAvidin

Transfected HEK 293T cells (two 10-cm plates per condition; 5 μg DNA/plate) were detached using TrypLE (1 mL/plate), quenched with media (4 mL/plate), washed 3X with cold 1X PBS pH 7.4 (10 mL), and centrifuged at 300 x g for 10 mins at 4 °C to pellet. The cell pellet was re- suspended in 300 µL of hypotonic buffer (10 mM KCl, 1.5 mM MgCl2, 10 mM HEPES pH 7.5, 1X protease inhibitor (PI) cocktail (Roche cOmplete Mini EDTA-free cocktail)) and incubated on ice for 5 min. Samples were centrifuged at 300 x g for 10 minutes at 4 °C to pellet the nuclei. After removal of supernatant, nuclei were re-suspended in 450 µL labeling buffer (25 mM HEPES pH 7.5, 50 mM NaCl, 5 mM MgCl2, 1% NP40, 1 mM TCEP, and 1X PI cocktail. 5-Bn- DTB-NAD^+^ (1 mM in 10%; 50 µL) was added for 100 μM 5-Bn-DTB-NAD+ and 1% DMSO final concentration). Next, samples were incubated at 30 °C for 2 h at 400 RPM and vortexed in ∼15-minute intervals to re-suspend nuclei that settled down during incubation. The reaction was stopped by the addition of PARP inhibitor Phthal01 (19). Samples were precipitated in 9 mL of MeOH/CHCl3/H2O (4:1:4) and centrifuged at 5000 x g for 10 min at 4 °C. The central protein disc at the interface of the two layers was removed and washed with MeOH (1.5 mL), MeOH/CHCl3 (0.9 mL; 2:1), re-pelleted as before, and air dried. Pellets were re-suspended in 500 µL 6 M urea in 1X PBS pH 7.4 with 0.5% SDS at 37 °C for 60 min. Samples (300 µL at 2 mg/mL) were added to 100 µL pre-equilibrated Pierce™ High Capacity NeutrAvidin™ Agarose (Thermo Scientific™ 29202) (washed once with 6 M urea in 1X PBS pH 7.4 with 0.5% SDS) and rotated end-over-end for 2 h at room temperature. Samples were centrifuged at 4000 x g for 10 min. The supernatant was removed, and the resin was washed 1X with 6 M urea in 1X PBS pH 7.4 with 0.5% SDS (300 µL), 3X with 0.2% SDS in 1X PBS pH 7.4 (300 µL), and 3X with 1X PBS pH 7.4 (300 µL). Samples were eluted with 4% SDS with 5 mM biotin in 1X PBS pH 7.4 (100 µL) at 37 °C for 1 hr. All samples were boiled at 95 °C for 5 min, analyzed via SDS-PAGE, and transferred for WB.

### Generation of GFP-WT-PARP7 and GFP-ΔKBR3-PARP7 doxycycline-inducible CT26 cell lines

Lentiviruses for expression of GFP-WT-PARP7 and GFP-ΔKBR3-PARP7 were generated in HEK 293T cells as described previously (87). Briefly, HEK 293T cells were co-transfected with pCW57-GFP-WT-PARP7 or pCW57-GFP-ΔKBR3-PARP7 and 3rd generation lentiviral packaging plasmids (pLP1, pLP2, pLP-VSVG, Invitrogen) using calcium phosphate (CalPhos) transfection (Clontech). 24 h and 48 h after transfection, lentiviruses were collected, filtered (45 mm, cellulose acetate), and concentrated using ultracentrifugation (37,000 x g, 2 h, 4°C).

Approximate lentiviral titers were determined using Lenti-X™ GoStix™ (Clontech). Concentrated lentivirus was used to transduce CT26 PARP7 KO cells (Synthego), at ∼80% density. 72 h post-transduction, cells were split and selected with 25 µg/ml blasticidin. Cells were maintained under blasticidin selection.

### Immunoblotting

For immunoblotting, cells were harvested after a wash with ice-cold PBS and stored at −80°C until lysis. Lysis was carried out with cell lysis buffer (50 mM HEPES pH 7.4, 150 mM NaCl, 1 mM MgCl2, 1% Triton X-100) supplemented with 100 μM TCEP, 10 μM Trichostatin A (TSA), 1X phosphatase inhibitors (Sigma cocktail 2&3), and 30 μM Pan-PARP inhibitor Pthtal01 (19) Lysates were quickly vortexed and clarified by centrifugation at 14,000 x g at 4°C for 10 min. Protein concentration from each lysate was determined using the Bradford assay (Bio- Rad) to normalize protein loading. 4X sample buffer (40% glycerol, 200 mM Tris-HCl (pH 6.8), 8% SDS, 4% β-mercaptoethanol, 0.02% bromophenol blue) was added to lysates, and samples were boiled at 95°C for 5 min and loaded to SDS-PAGE. Gels were transferred to nitrocellulose membrane using Trans-Blot Turbo Transfer System (BioRad). Membranes were blocked with 5% milk (Carnation) in TBS (10 mM Tris-HCl pH 7.5, 150 mM NaCl) containing 0.1% Tween 20 (TBST). The membranes were then incubated with the respective antibodies.

To achieve complete solubilization of cellular proteins, including nuclear and chromatin-associated proteins, for stability assays and evaluation of PARP7 auto-MARylation, cells were lysed in 50 mM HEPES pH 7.4, 150 mM NaCl, 1 mM MgCl_2_, 1% SDS and 300 U/ml benzonase (Millipore Sigma, 70664) at room temperature. After quantification, dilution, and addition of 4X SDS-sample buffer, samples were processed as described above.

### Co-Immunoprecipitations

Cells were collected, washed with ice-cold PBS, and stored at −80°C until lysis. Cells were lysed in lysis buffer (150 mM NaCl, 25 mM Tris, 1% Nonidet P-40, 10% glycerol, freshly supplemented with 1X phosphatase inhibitors (Sigma cocktail 2&3), and 30 μM Pan-PARP inhibitor Pthtal01). 1 mg of each lysate was incubated with 15 μL of GFP-Trap® magnetic agarose beads (ChromoTek) *for one hour at* 4°C *with agitation. Beads were pulled down with a magnetic base and washed four times with 0.5 mL of low salt washing buffer (*50 mM NaCl, 25 mM Tris, 0.1% Nonidet P- 40) to evaluate PARP7-partners interaction and high salt washing buffer (500 mM NaCl, 25 mM Tris, 0.1% Nonidet P-40) to evaluate CBP and p300 ADP-ribosylation. Beads were eluted by adding 1X sample buffer and heated at 95°C for 5 min. Western blotting was carried out as described above.

### Cellular fractionation

Cells were harvested and immediately lysed with cytoplasmic extraction buffer (10 mM HEPES pH 7.5, 10 mM KCl, 0.1 mM EDTA) supplemented with freshly added 0.4% NP-40, 1 μM DTT, 10 μM Trichostatin A (TSA), 1X phosphatase inhibitors (Sigma cocktail 2&3), and 30 μM Pthtal01. Cells were scraped, re-suspended by gently pipetting up and down, and transferred to a pre-chilled tube. Cell preparations were centrifuged at 14,000 x g for 3 min. Supernatants were saved. Pellets were resuspended in nuclear extraction buffer (10 mM HEPES, pH 7.5, 400 mM NaCl, 1 mM EDTA, 10% Glycerol), supplemented with the inhibitors above. Tubes were vortexed for 15 s and placed in a rotator for 2 h, vortexing at the 1 h mark. After 2 h, tubes were vortexed for 15 s and centrifuged at 14,000 X g for 5 min. Protein concentration was calculated as described above.

### ELISA assays

Cells were seeded into 6-well plates with RPMI containing 1X pyruvate and 10% Tet-free FBS (Gibco A4736301). After 24 h, cells were treated with doxycycline. Media was harvested after 20 h, and IFNβ was measured using the Verikine ELISA kit according to the manufacturer’s instructions. Cells were lysed and analyzed by immunoblot.

Immunofluorescence imaging

### In vitro calf Lambda phosphatase and USP21 assay

CT26 cells were treated with DMSO or RBN2397 for 24 h before stimulation with DMXAA. Immediately after harvest, nuclei were fractionated as described before and diluted 1:3 in dilution buffer (20 mM HEPES pH 7.5, 10% Glycerol, with freshly added 0.4% NP-40, 1 μM DTT, 10 μM Trichostatin A (TSA). Each diluted nuclei preparation was separated into three tubes containing 1X NEB-Buffer for protein metallo phosphatases and 1X MnCl2 (1 mM). The reaction began with the addition of 400 U of λPP, 5 μM USP21, or water to each tube. Mixtures were incubated at 37 °C for 30 min. The reactions were stopped by the addition of 4X SDS sample buffer, boiled for 5 min at 95 °C, and run in 6% SDS-PAGE. Changes in the migration of p300 were detected by Western blot.

### Immunofluorescence imaging

HEK 293T cells were plated on PDL-coated (Cultrex) cover slips (13 mm, No 1.5, Fisher) transfected with GFP-WT- or GFP-ΔKBR3-PARP7 as described above. GFP-WT-PARP7 cells were treated with dCBP-1 (1 mM) for 6 h. The media was removed, and the cells were fixed using 4% PFA in PBS (15 min, RT). Cells were washed with PBS (3 x 5 min), permeabilized with 0.2% TX-100 (5 min, RT), and incubated with Hoechst 33342 (Invitrogen) in PBS (10 min, RT). Cells were washed with PBS (3 x 5 min), rinsed in ddH_2_O, and mounted on glass slides using ProLong™ Gold Antifade Mountant (Invitrogen).

### GST-PARP7 expression and purification

GST-PARP7 plasmid was transformed into BL21(DE3) competent cells and grown in Terrific Broth media supplemented with ampicillin. Induction was carried out at 0.6–0.8 OD_600_ using 0.5 mM IPTG, and cells were allowed to grow overnight at 16°C. Bacterial pellet was lysed in lysis buffer (20 mM Tris-HCl pH 8.0, 500 mM NaCl, 1% Triton-X, 10% Glycerol, 0.1 mM ZnCl2, 2 mM DTT, Sigmafast™ protease inhibitor (S8830-20TAB). The cleared and filtered lysate was then incubated with pre-washed Glutathione Sepharose 4B agarose resin for 1 hour. Resin was washed with lysis buffer and 1M NaCl wash buffer (20 mM Tris-HCl pH 8.0, 1 M NaCl, 1% Triton-X, 10% Glycerol, 0.1 mM ZnCl_2_, 2 mM DTT). GST-PARP7 was eluted in 100 mM Tris-HCl pH 8.0, 150 mM NaCl, 10% Glycerol, 0.1 mM ZnCl2, 40 mM glutathione. Purified GST-PARP7 was stored at −80 °C.

### AlphaFold modeling

The predicted hPARP7 structure was obtained from the AlphaFold Protein Structure Database (alphafold.ebi.ac.uk). Interaction studies were then performed using ColabFold. Structures were analyzed on Chimera-X.

### Statistical Analysis

The experiment data were analyzed by Student’s *t*-test or one-way ANOVA in GraphPad Prism 8 software. *p* < 0.05 was considered statistically significant.

## Supporting information

SupFigures

## Acknowledgments

The authors thank both current and former Cohen members for their valuable discussions on experimental design, data analysis, and interpretation, as well as for providing general advice and feedback on this project. We thank Dr. Andreas Marquardt and Dr. Anna Sladewska-Marquardt of the Proteomics Center at the University of Konstanz for their assistance with the mass spectrometric experiments. Dr. Jon Tullis is the recipient of the Jane Coffin Childs postdoctoral fellowship. This work was supported by the National Institutes of Neurological Disorders and Stroke 2R01NS088629 (to MSC).

## Funding

National Institutes of Neurological Disorders and Stroke 2R01NS088629 (PV, MSC).

## Author contributions

Each author’s contribution(s) to the paper should be listed (we suggest following the CRediT model with each CRediT role given its own line. No punctuation in the initials.

Conceptualization: IRS, SR, AM, MSC

Methodology: IRS, SR, DJ, JO, JT, MSC

Investigation: IRS, SR, RM, DJ, JT, JO

Visualization: IRS, SR MSC

Supervision: MSC

Writing—original draft: IRS, MSC

Writing—review & editing: IRS, JT, MSC

## Competing interests

The authors declare no competing interests.

## Data and materials availability

All data are available in the main text or the supplementary materials. Materials, including plasmids and compounds, are available upon request.

## Notes

### Competing Interest Statement

The authors have declared no competing interest.

