## Supplementary material for "The IFN I response in tumor cells is shaped by PARP7–p300/CBP interactions through distinct loss- and gain-of-function mechanisms": SupFigures

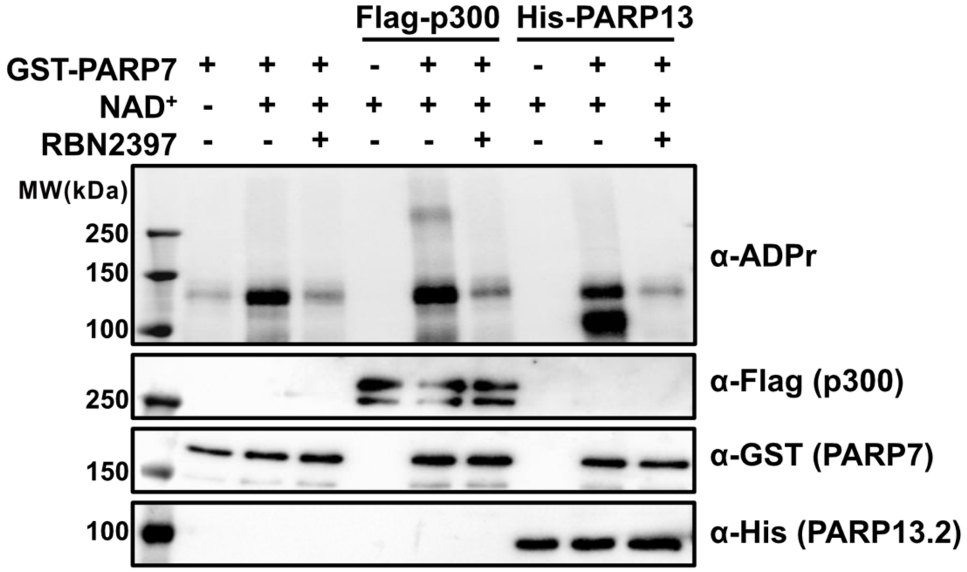


**Fig. S1. PARP7 MARylates p300 *in vitro*.** 100 nM of recombinant Flag-p300 were incubated with 100 nM GST-PARP7 in the presence and absence of 0.1 mM $NAD^+$ for 1 h at 30 °C. His-PARP13.2 was used as a positive control, and 300 μM RBN2397 was used to inhibit PARP7-mediated MARylation.

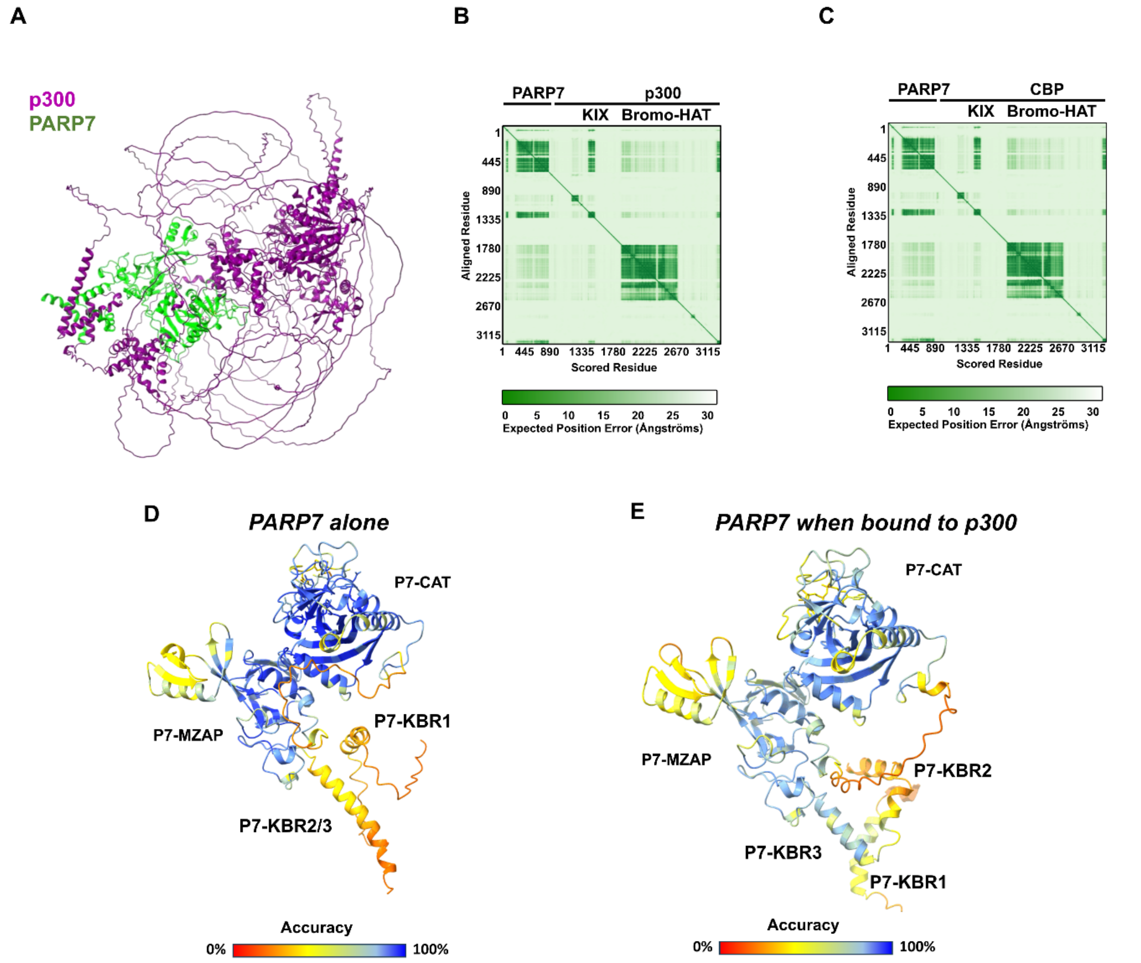


Fig.

**Fig S2. High accuracy prediction of PARP7-p300/CBP interaction.**
(**A**) AlphaFold 3 prediction of the full length PARP7 (green) in complex with p300 (purple).
(**B**) Predicted aligned error (PAE) plot of PARP7-P300 complex.
(**C**) Predicted aligned error (PAE) plot of PARP7-CBP complex.
(**D**) Predicted PARP7 structure in the AlphaFold database (AF-Q7Z3E1-F1-v4).
(**E**) AlphaFold 3 prediction of interaction between PARP7 and the KIX domain of p300.

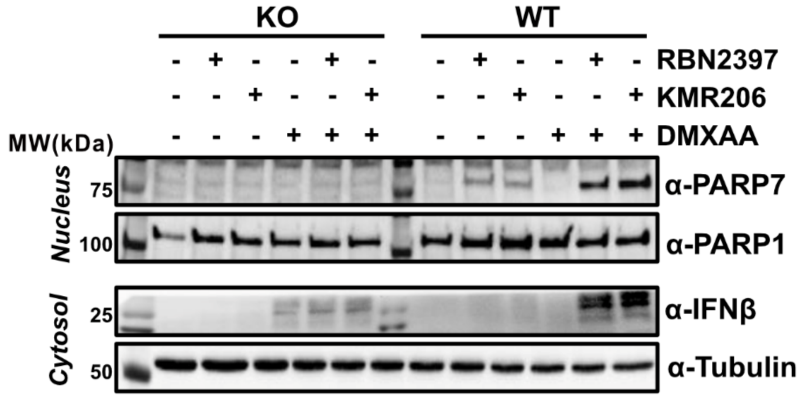


Fig. S3. Inhibitor-induced gain-of-function effect on IFNβ is dependent on PARP7 expression. Parent and PARP7 KO CT26 cells were incubated with either RBN2397 or KMR206 for 20 h and treated with ± 10 μg/mL DMXAA for 6 h. Cytoplasmic and nuclear fractions were se
